# Non-lethal exposure to H_2_O_2_ boosts bacterial survival and evolvability against oxidative stress

**DOI:** 10.1101/575134

**Authors:** Alexandro Rodríguez-Rojas, Joshua Jay Kim, Paul Johnston, Olga Makarova, Murat Eravci, Christoph Weise, Regine Hengge, Jens Rolff

## Abstract

Unicellular organisms have the prevalent challenge to survive under oxidative stress of reactive oxygen species (ROS) such as hydrogen peroxide (H_2_O_2_). ROS are present as by-products of photosynthesis and aerobic respiration. These reactive species are even employed by multicellular organisms as potent weapons against microbes. Although bacterial defences against lethal and sub-lethal oxidative stress have been studied in model bacteria, the role of fluctuating H_2_O_2_ concentrations remains unexplored. It is known that sub-lethal exposure of *Escherichia coli* to H_2_O_2_ results in enhanced survival upon subsequent exposure. Here we investigate the priming response to H_2_O_2_ at physiological concentrations. The basis and the duration of the response (memory) were also determined by time-lapse quantitative proteomics. We found that a low level of H_2_O_2_ induced several scavenging enzymes showing a long half-life, subsequently protecting cells from future exposure. We then asked if the phenotypic resistance against H_2_O_2_ alters the evolution of resistance against oxygen stress. Experimental evolution of H_2_O_2_ resistance revealed faster evolution and higher levels of resistance in primed cells. Several mutations were found to be associated with resistance in evolved populations affecting different loci but, counterintuitively, none of them was directly associated with scavenging systems. Our results have important implications for host colonisation and infections where microbes often encounter reactive oxygen species in gradients.

## I. Introduction

The ability to elicit a stress response when encountering repeated stress relies on ‘remembering’ a similar event from the past (memory), a trait common to many biological entities [1]. During the course of an infection or the colonisation of a host, bacteria encounter increasing and repeated stress imposed by the host immune system [2,3]. Obata *et al*., for example, recently demonstrated that low levels of H_2_O_2_ in the gut of *Drosophila melanogaster* shape the composition of the gut microbiota resulting in differential survival of the flies [3]. Here, we report how bacteria respond to the exposure to low levels of reactive oxygen species (ROS) and how this impacts bacterial fitness. We then investigate if the phenotypic response to a sub-lethal dose of H_2_O_2_ facilitates resistance evolution, hence providing a test of the plasticity-first hypothesis, which proposes that environmentally initiated phenotypic change precedes or even facilitates evolutionary adaptation [2,4].

In *Escherichia coli*, the defences against oxidative stress depend on transcriptional regulators such as OxyR or SoxR that detect changes in the redox balance. They also induce the production of detoxifying enzymes, DNA repair and protection systems and other proteins [5]. Oxidation of OxyR by H_2_O_2_ leads to the formation of an intramolecular disulfide bond between cysteine residues 199 and 208. Oxidized OxyR positively regulates catalases and peroxidases. OxyR is deactivated by enzymatic reduction with glutaredoxin I (Grx) or thioredoxin (Trx). Because the Grx/GorA system is itself transcriptionally regulated by OxyR, the whole response is self-regulated in a homeostatic feedback loop [6].

The OxyR-mediated oxidative stress response results in scavenging of H_2_O_2_ and mitigates the toxicity of this by-product of aerobic metabolism. It includes the induction of Suf proteins that form a complex to supply apo-enzymes with iron-sulfur clusters. The Suf system replaces the normal iron-sulfur cluster supply system (Isc), required for critical biochemical pathways such as respiration, which is disrupted by H_2_O_2_ stress [7,8]. The iron-sulphur clusters of dehydratases are one of the most H_2_O_2_-sensitive systems. The repair of those clusters by Suf is necessary to prevent the failure of the TCA cycle. The iron-sulfur clusters from enzymes that employ ferrous iron as a co-factor can increase the risk of fuelling a Fenton reaction. Active OxyR also induces Dps, a ferritin-class protein, that strongly suppresses the amount of DNA damage by sequestering the unincorporated iron [9]. OxyR mutants accumulate ROS at much higher levels than the wild-type strain during growth even in the absence of H_2_O_2_ which also accounts for its high sensitivity [10].

The spontaneous reaction of H_2_O_2_ with free ferrous iron (Fe^2+^) at physiological pH, oxidising iron to Fe^3+^ and generating hydroxyl radicals and water, is named the Fenton reaction. Hydroxyl is a strong non-selective radical that damages many cellular components, particularly DNA [11,12]. H_2_O_2_ impedes the function of the Fur regulatory protein and can directly damage many cell components but is less toxic than other reactive oxygen species such as hydroxyl. These radicals are responsible for DNA damage, indirectly promoted by H_2_O_2_ as a consequence of the Fenton reaction [13].

A classic paper by Imlay *et al*. describes that the pre-treatment of *E. coli* with a low dose (60 µM) of H_2_O_2_ can increase the survival upon subsequent exposure to an otherwise lethal dose (30 mM) [14]. However, neither the duration of such priming responses nor the molecular mechanisms of its maintenance, i.e. the memory, have been studied. Our study has two main aims. First, we investigated the main factors in the H_2_O_2_ priming response and for how long the response is sustained. Second, we used this system to test the hypothesis that inducible phenotypes accelerate adaptive evolution [4]. Therefore, we experimentally evolved *Escherichia coli* under increasing concentrations of H_2_O_2_ with and without priming. For H_2_O_2_-resistant populations evolved after priming and non-priming regimes, genome re-sequencing analyses were performed to identify mutations. We focus on growing bacteria (exponential phase), as this better represents infection or colonisation of host surfaces including the gut [2,3]. Moreover, this allows us to focus on the priming response in proliferating cells, where the response to H_2_O_2_ differs from that in stationary phase. While the response to the H_2_O_2_ in exponentially growing bacteria is mostly controlled by OxyR, RpoS, the master regulator of the general stress response controls a pronounced phenotypic H_2_O_2_ tolerance during stationary phase [15,16].

## II. Results and Discussion

### Priming by H_2_O_2_ results in higher bacterial survival

We found that, in our conditions, the minimal inhibitory concentration for H_2_O_2_ is 1 mM. This concentration was subsequently used as a reference for 30-minute killing curves that show a clear dose-effect in survival rate (Fig 1a, ranging from 50 µM to 1 mM, Fig 1a, *p*=2.1×10^−16^, DRC model fitted based on maximum likelihood [17]). Based on these results, we primed cells for 30 min with 0.1 mM H_2_O_2_ and challenge the cultures with 1 mM H_2_O_2_ 90 min after priming stimulus. The priming concentration (0.1 mM) is the maximum dose that shows no difference in growth rate for each time-point compared to non-treated cells (S1 Fig, S1 Table). Primed (pre-treated) populations of *E. coli* showed higher survival than naïve cells by more than one order of magnitude (Cox proportional hazard model, *p*<0.05, Fig 1b). We also determined that the priming response contributes to a more efficient removal of H_2_O_2_ by quantifying it in the supernatant of the cultures (S2 Table). After 15 minutes, H_2_O_2_ dropped from 1 mM to 0.723 mM for control while in pre-treated cultures H_2_O_2_ decreased to 0.140 mM (*p*=0.00014, pre-treated versus control). For 30 minutes, the level of H_2_O_2_ went down to 0.377 mM for control cultures while in pre-treated cultures, it decreased to 0.0222 mM (*p*=0.00035).

**Fig 1.**
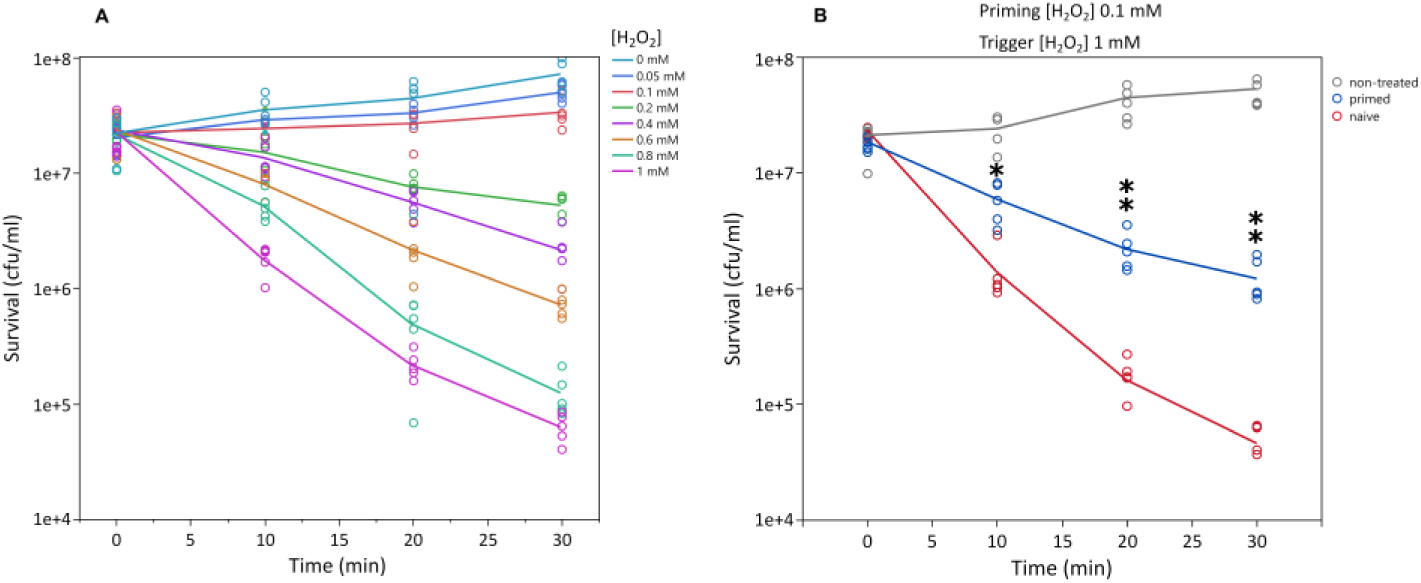
Hydrogen peroxide killing curves of *E. coli* MG1655. Panel A shows bacterial sensitivity to 1 mM H_2_O_2_ with a strong dose-response effect (*p*=2.1×10^−16^, DRC model fitted based on maximum likelihood). Panel B shows the priming effect or improved response when cells are treated in advance with a non-lethal dose (0.1 mM). Curves represent the mean of five independent cultures per time point. Asterisks represent significant differences (Welch’s test, one asterisk for *p*<0.05 and two asterisks for *p*<0.01). Only comparisons between primed and non-primed groups are shown.

Although there is no difference in growth rate, priming with 0.1 mM H_2_O_2_ imposes a small cost (approximately 4%, S1 Fig, S1 Table). The cost can be detected by comparing the empirical area under the curve (p=0.0002, 0.9619 fold-change) or the carrying capacity of the curve (*p=*0.0001, 0.9565 fold-change) from primed bacteria culture versus non-treated control by allowing bacteria to grow until the stationary phase. No other parameter such as initial population size, growth rate, or doubling time is affected. Therefore, differential survival can be attributed only to cell response but not to cell growth arrest. The cost can be explained by even very low concentrations of H_2_O_2_ damaging the iron-sulfur clusters, thereby compromising respiration. However, at low bacteria density in rich medium, *E. coli* can grow very fast by fermentation [18]. This wasteful strategy allows quick generation of ATP at the expense of the medium and can explain the small difference in carrying capacity and the area under the curve of the growth curves (Fig S1, S1 Table). In our experiments, this cost has no consequences because bacteria are maintained in low density and in exponential growth. This scenario should be similar while starting an infection or colonisation of a host’s gut.

These results are in agreement with data previously reported by Imlay *et al*. [11,14], showing survival protection even to doses as high as 30 mM, a concentration close to H_2_O_2_ usage as a disinfectant. Our concentrations are also in the range of some *in vivo* situations. For example, tailfin transection on zebrafish larvae induces a rapid increase in H_2_O_2_ levels ranging from 100–200 μM in the wound margins [2]. In some cases, more than 100 μM of H_2_O_2_ have been reported in human and animal eye vitreous humour and aqueous humour [19]. In plants, the average tissue concentration of H_2_O_2_ is around 1 mM, but under stress it can as high as 10 mM [20]. This could be relevant for foodborne pathogens transmitted by consumption of contaminated vegetables.

### Duration of the priming response

As the priming response protects the cells effectively from an otherwise lethal exposure to H_2_O_2_, an important question is for how long this response remains effective. To address this, we pre-treated *E. coli* again with 0.1 mM H_2_O_2_ for 30 minutes, but we applied the higher dose (1 mM, trigger of the priming response) at different time-points (30, 60, 90, 120 and 150 minutes after the 30 min priming period, cell density kept constant by appropriate dilution). We observed a significant decay of the priming response from 120 minutes after H_2_O_2_ pre-treatment removal (Fig 2), approximately four divisions, suggesting that the priming effect is also trans-generational. After 150 minutes, the survival rate of primed populations no longer differed from naïve populations. Another study has shown long-term memory based on an epigenetic switch that controls a bimodal virulence alternation also in the scale of hours in *E. coli* [21].

**Fig 2.**
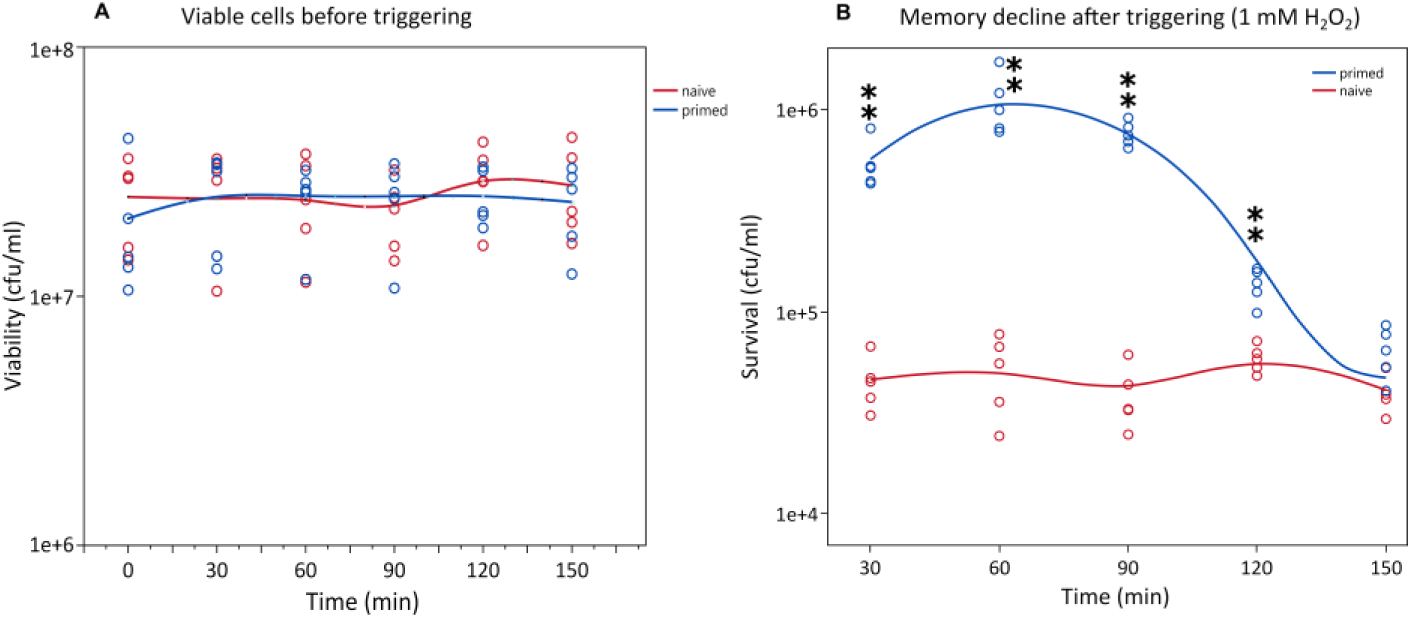
Bacterial survival from naïve and primed populations triggering at 30 minute intervals after the priming stimulus (memory of the priming response). During memory decline experiment, the viability of the cells remains unaltered before the addition of the trigger (A). The priming memory declined over time after induction of priming response with 0.1 mM H_2_O_2_ (trigger) (B). Asterisks indicate significant differences between each time-point pair (Welch’s test, one asterisk for *p*<0.05 and two asterisks for *p*<0.01).

How is the state of priming maintained for up to five generations? Bacteria can store information about recent stress via stable transcripts or proteins [21,22]. We investigated the memory at the protein level and initially studied the impact of priming and trigger concentrations (0.1 and 1 mM) on the proteome of *E. coli* by quantitative LC-mass spectrometry. We detected upregulation of many of the known enzymes that are induced by H_2_O_2_ just 5 minutes after the addition of H_2_O_2_ (Fig 3, and S3 Table and S4 Table).

**Fig 3.**
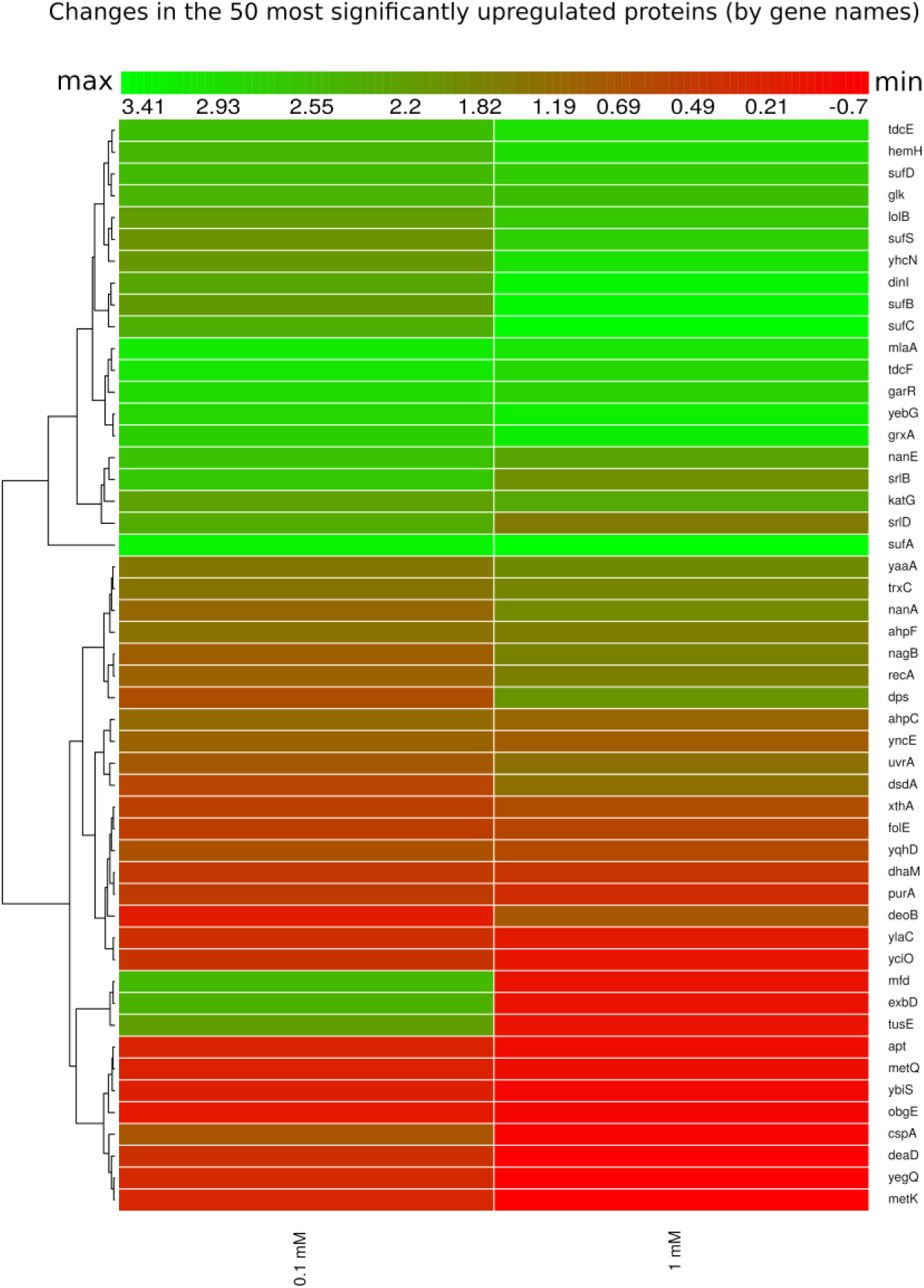
Heatmap of relative protein expression based on label-free quantification by liquid chromatography/mass spectrometry (LC-MS). Hierarchical clustering of the intensities was performed using Euclidean distances between means. Rows indicate the fluctuation of protein (by gene names) level at 5 minutes after addition of 0.1 and 1 mM H_2_O_2_. Intensity ranges of the log2 fold-changes are given from highest intensity (green) to lowest (red). Only the 50 most statistically significant up-regulated proteins are shown, taking as a reference the 0.1 mM concentration.

For both concentrations used, we detected and quantified many proteins belonging to the OxyR regulon. Furthermore, many other genes, such as *ahpC*/*F, xthA* and *suf* operons showed a weak difference or no response at all. The samples in this experiment were taken after only 5 minutes of treatment. The rationale behind this design is that the availability of viable cells after 1 mM H_2_O_2_ treatment would be too low at a later stage. At a concentration of 0.1 mM, however, it is possible to collect viable cells for any other time-point after exposure. These results are presented in Fig 4.

**Fig 4.**
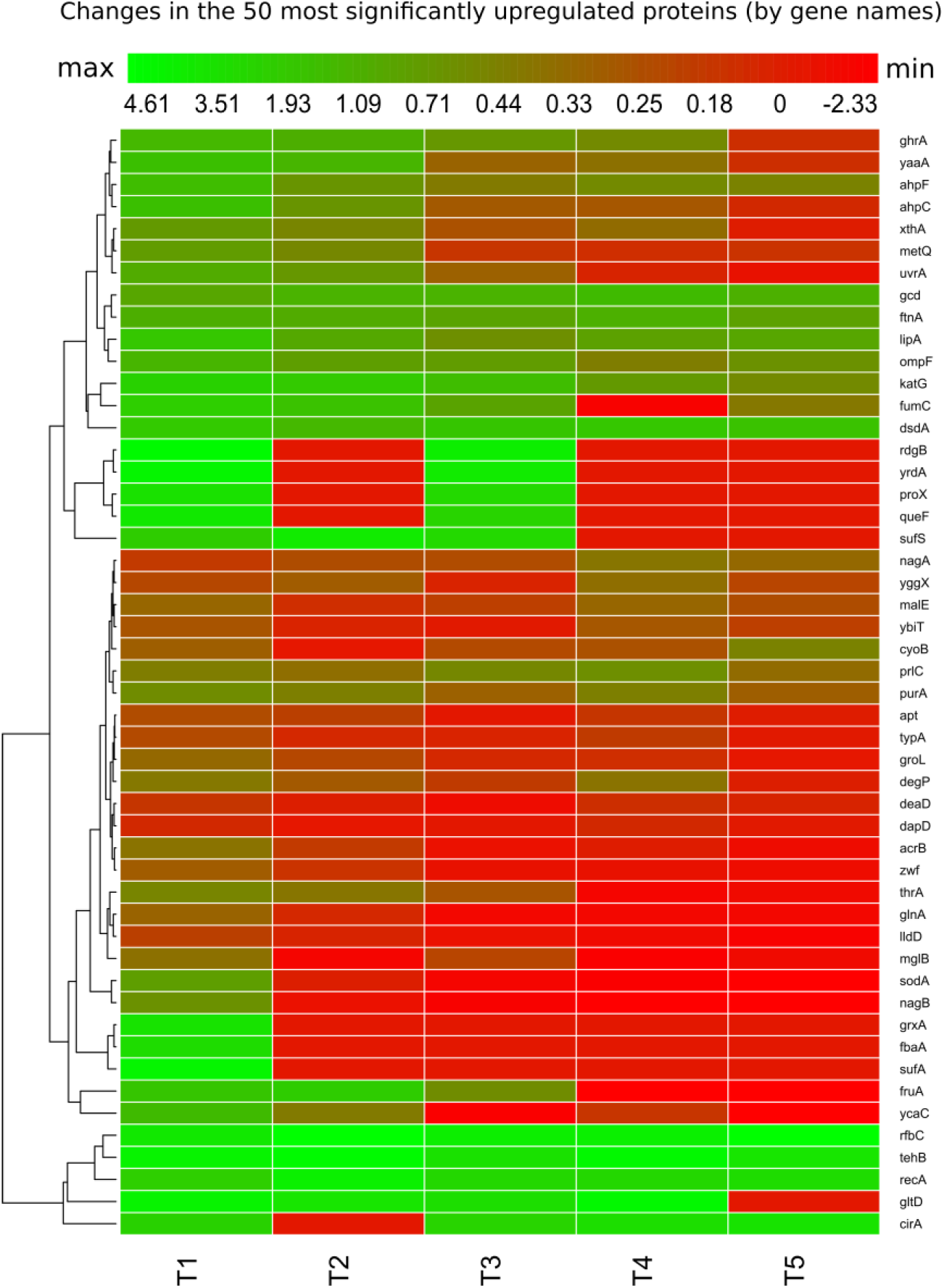
Heatmap of relative protein expression based on quantitative label-free quantification by liquid chromatography/mass spectrometry (LC-MS). Hierarchical clustering of the intensities was performed by using Euclidean distances between means. Rows indicate the fluctuation of protein (by gene names) level at different time points. Intensity ranges of the log2 fold-changes are given from highest intensity (green) to lowest (red). Only the most 50 statistically significant up-regulated proteins at time point T1 (30 minutes after priming) are represented to follow their fluctuation in the other time points with 30 minutes between them.

To explore the temporal dynamics of the proteome after a 30-minute stimulus with H_2_O_2_ (0.1 mM), we followed the changes over almost 3 hours (five time points: 30, 60, 90, 120 and 150 minutes). The decline in anti-H_2_O_2_ protein levels correlates well with the decrease of the response (Fig 4). Proteins such as KatG, AhpF or RecA declined slowly after removal of the H_2_O_2_, consistent with a sustained production with a minor contribution of dilution due to cell division and slow degradation rate. These results indicate that many of these proteins are stable and show a significantly higher abundance than in the control even at 150 minutes after H_2_O_2_ treatment. Other initially induced proteins such as GrxA, YaaA and XthA, SufA, SufS, AcrA-AcrB had completely declined at this point indicating that these proteins may be subject to proteolysis and have shorter half-lives (Fig 4) but also that the stressful situation is alleviated. The overall results of this proteomic experiment show that the memory of the priming response in *E. coli* is mediated by the scavenging proteins such as KatG and AhpCF. The primary amino-acid sequence is informative about the *in vivo* half-life a protein. An N-end rule-based prediction of the half-life for some of the proteins that are responsible for the memory additionally is consistent with our findings obtained by the proteomic approach (S5 Table).

We visualised the global impact of H_2_O_2_ on bacterial physiology using a network analysis based on protein-protein interactions and function [23]. This network analysis provides information on protein level alterations. It integrates protein-protein interactions, including indirect (functional) and direct (physical) associations [23]. We have projected our proteomic datasets over the established interactions of *E. coli* proteins to illustrate the scope of the priming response, including toxic effects. The proteome response to the priming concentration (0.1 mM H_2_O_2_ during 30 minutes), resulted in a high degree of connectivity of protein-protein interactions and functional relation of both, up- and down-regulated proteins (Fig 5). If we compare our network with a large-scale protein-protein interaction network of *E. coli* [24], we find a wide perturbation including the most important nodes. This analysis also points to proteome-wide readjustments to cope with H_2_O_2_ stress and shows the profound impact of oxidative stress across the entire proteome even at a low dose that does not change the growth rate in a rich medium. The damage induced to iron-sulfur clusters even at a low H_2_O_2_ concentration (0.1 mM) might have no detectable impact on bacterial cell growth in early exponential phase, since under these conditions most of the energy is obtained by fermentation, as shown for *E. coli* previously [18]. This means that at low cell density in rich medium, the consumption of the resources does not drastically change the medium properties and it is advantageous for bacteria to use a costly strategy that provides them with fast energy via fermentation which supports faster growth compared to aerobic respiration.

**Fig 5.**
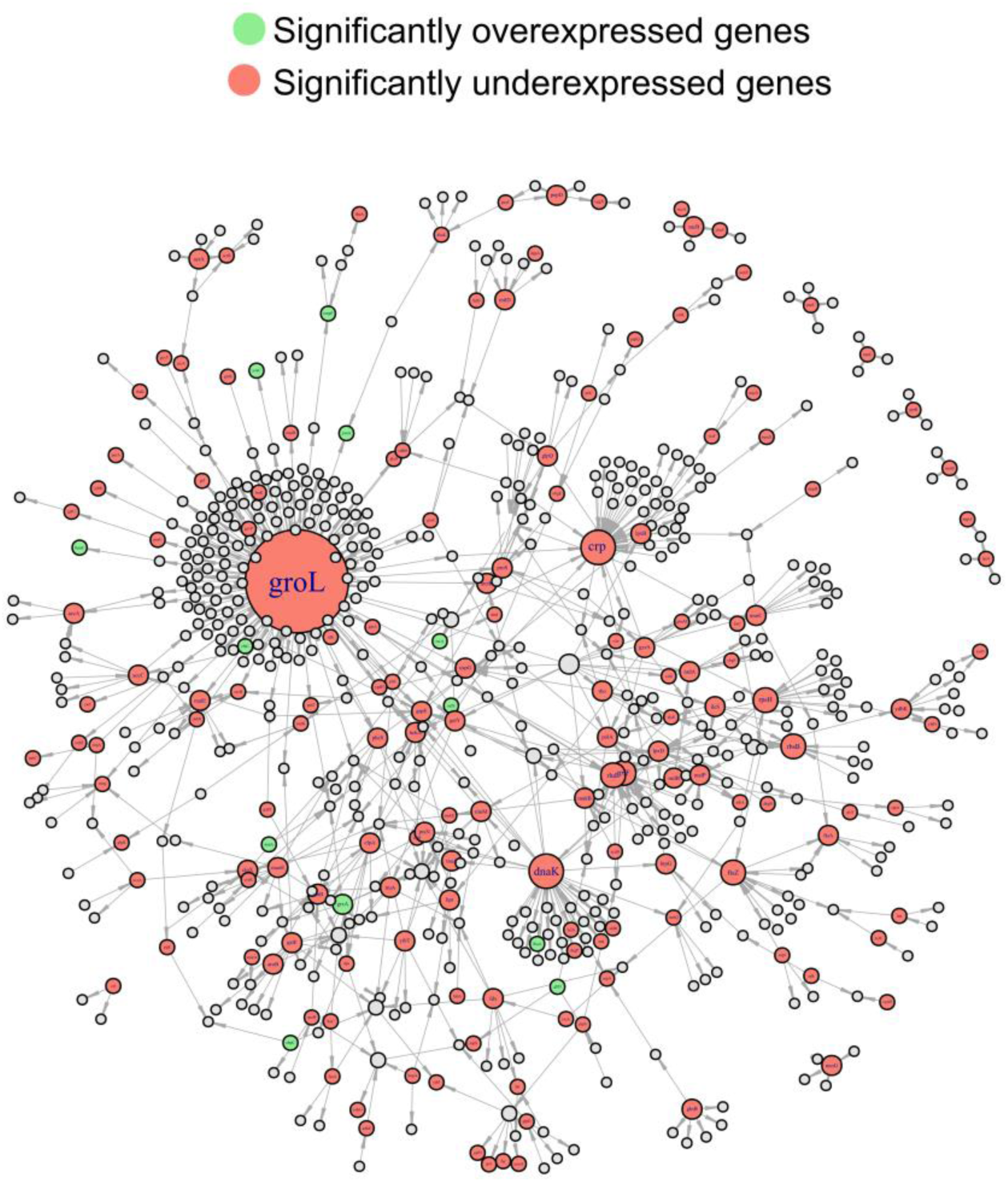
Projection of differentially translated proteins 30 minutes after addition of H_2_O_2_ on protein-protein Interaction network of *E. coli* K12. Pale green nodes indicate up-regulated proteins while pale red ones represent down-regulated ones. Note that the most critical nodes (DnaK and GloL) of the protein-protein interaction network are affected. The interaction among nodes shows the proteome-wide impact of H_2_O_2_.

The memory could also be based on long-lived transcripts. Therefore, we sequenced the full transcriptome after exposure to H_2_O_2_. RNAseq captured both sRNA and mRNA and we sampled just before the decline of the response (120 minutes after removal the stimulus). The transcript with the greatest induction was OxyS, a small regulatory RNA (sRNA) induced by active OxyR (S6 Table and S7 Table). OxyS regulates several genes, but although several targets have been identified, its function is not fully understood [25]. We did not detect upregulation of other transcripts under OxyR regulation.

OxyS is a potential candidate to explain the duration of priming because it is relatively stable with a half-life between 18 to 20 minutes during the exponential phase [25–27]. To explore this possibility, we used an *oxyS-*deficient mutant to test its sensitivity to H_2_O_2_ over 30 minutes. We did not find any significant differences in sensitivity (S2 Fig) consistent with prior reports [26–28]. Although OxyS did not provide us with a mechanism to explain the duration of the memory, its stability could play some role in alleviating DNA damage as recently suggested [29]. However, such protection does not seem to have a significant impact on cell survival.

Based on the comparison of proteomic and transcriptomic data it seems reasonable to assume that the capacity of the cells ‘to remember’ the stimulus is mainly based on the stability of scavenging proteins as documented in the proteomic dataset. These scavenging proteins remain present at higher concentrations than in the control samples as long as 120 minutes after removal of peroxide. This indicates that these enzymes are not degraded but probably their production continues after the stimulus during several cell divisions, which could explain the pattern that we observe in the decline of the response. It is important to consider that around 120 minutes, there were still significantly higher levels of KatG and AhpF than at the start of the experiment. Thus, the memory may require the contributions of additional genes whose relative expression decreased to low levels after two hours. It is also possible that specific enzymatic activity of KatG and AhpF gets lost over time, since the proteomic approach is based on protein identification by sequence, not by activity.

### Priming response is compromised by disrupting important H_2_O_2_-scavenging genes

The expression of genes important for H_2_O_2_ stress survival was also confirmed by qPCR 30 minutes after the addition of the chemical (0.1 mM). We found a significant up-regulation of selected genes such as *katG, ahpF*/*ahpC, dps, mntH* and *sufA* (S4 Fig and S8 Table). As previously described in the literature, *oxyR* and *fur* do not change in expression level when cells are treated with H_2_O_2,_ since their activation relies on switching between active or inactive forms of these proteins which depends on the intracellular level of H_2_O_2_ or iron respectively [6,7]. Many of these transcripts showed a strong induction at the RNA level (S8 Table). After 30 minutes, the level of induction of all genes showed a stronger response at the RNA level than at the protein level. In a normal situation, we would expect that a single molecule of RNA is translated into several proteins. This possibly indicates that under oxidative stress, the translation is inefficient and it is compensated by a high level of transcription probably due to damage of many cell components, including ribosomes as previously described [30].

To understand the priming response to H_2_O_2_ at the molecular level, we constructed a set of mutants for genes encoding key proteins identified in our proteomic dataset. In proliferating *E. coli*, OxyR is the major regulator controlling the cellular response to H_2_O_2_ [10]. We explored the involvement of OxyR in the priming response since many of the differentially expressed proteins were transcribed in an OxyR-dependent fashion. We found that by disrupting OxyR, there was a dramatic change in sensitivity to H_2_O_2_ with full loss of viability after 30 minutes (S2 Fig). The priming response is completely abolished (Fig 6), indicating that the enhanced survival, due to pre-exposure to H_2_O_2_, depends on the regulator OxyR. This regulator is a major transcription factor that protects *E. coli* against H_2_O_2_ during the exponential phase [10,31]. The active form of OxyR positively regulates dozens of genes.

**Fig 6.**
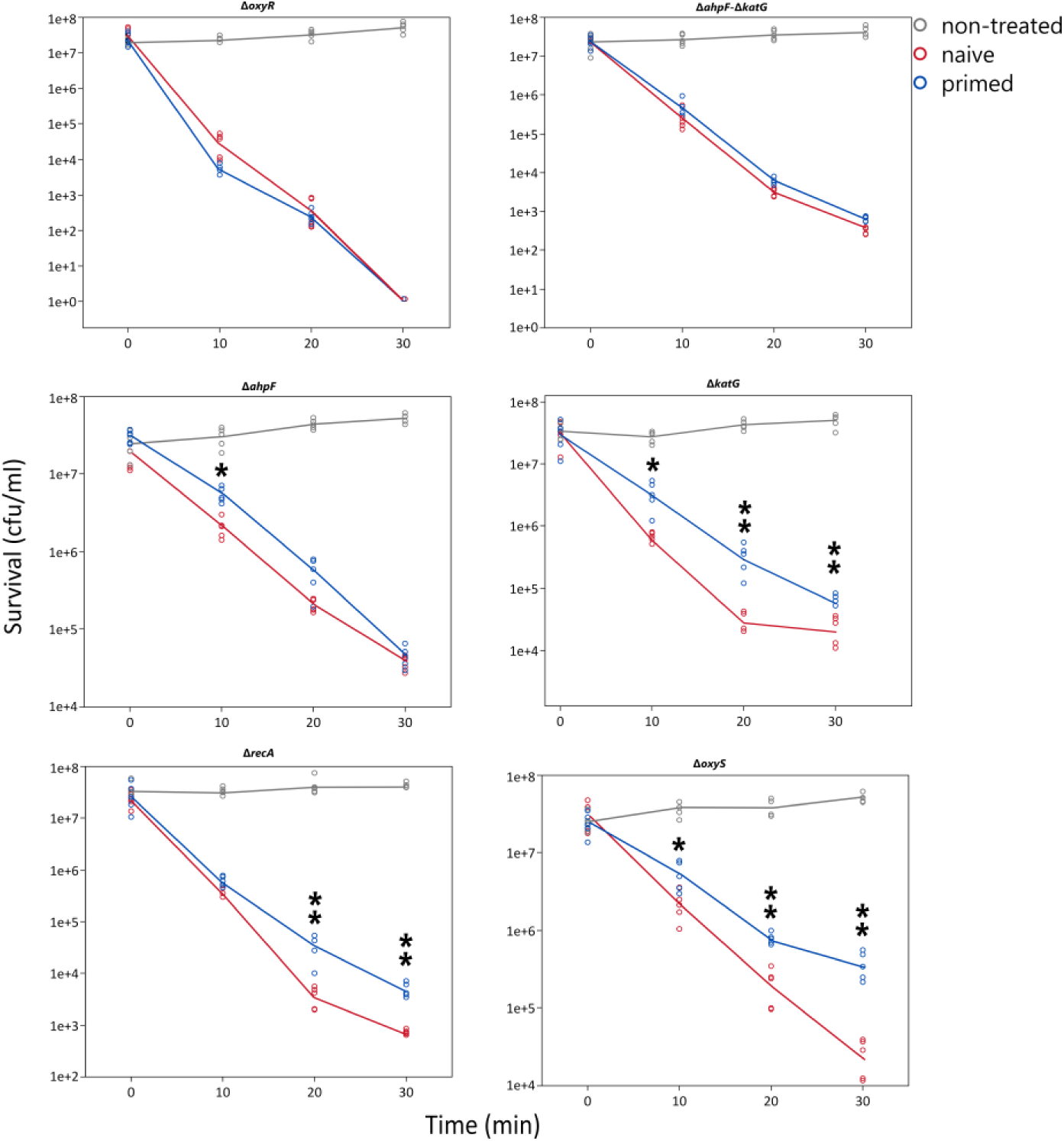
Priming experiments of different mutants of *E. coli* MG1655 in selected H_2_O_2-_responsive genes. Asterisks represent significant differences (Welch’s test, one asterisk for *p*<0.05 and two asterisks for *p*<0.01). Only comparison between primed and non-primed groups are shown.

Next, from the most highly expressed proteins and informed by published work [6,10], we generated a set of mutants that were used to determine the contribution of the respective genes to survival (Fig S2) and priming (Fig 6). Removal of KatG (catalase), AhpF (one subunit of the alkyl peroxidase AhpCF) or RecA (DNA repair) significantly decreased the survival in the presence of H_2_O_2_ (1 mM). The double mutant for KatG/AhpF showed even stronger sensitivity. However, the absence of other important proteins induced by H_2_O_2_ such as YaaA (decreases intracellular iron [31]), LipA (lipoate synthase, important for the repair of iron-sulfur clusters [32,33]), GltD (glutamate synthase subunit), GhxA or GhrA did not result in differential survival at 1 mM concentration. The OxyS sRNA was the transcript with the highest level of induction in the RNAseq data, but the mutant did not display any increased sensitivity to H_2_O_2_, it neither suppresses or decreases the priming response compared to the wild-type control. Assuming that H_2_O_2_ sensitivity and the priming response are closely linked, we included in the additional priming response experiment mainly those mutants with increased sensitivity (Fig 6 and S2 Fig). The removal of KatG indicates that catalase importantly contributes to the priming response, but it does not fully explain the protection observed for the WT strain (compare to Fig 1B). In the absence of AhpF, we also observe differences in priming response with the naïve state but not as pronounced as in the case of KatG-deficient strain. The double mutant defective in KatG and AhpF showed a dramatic decrease in the priming response but was still significantly different from naïve cells. Recently, a report showed that bacteria lacking the AhpF/C system suffer a severe post-stress recovery [34]. The absence of the AhpF/C system also contributes to the lethality of the mutants probably by the inability of the cell to cope with a low level of ROS after severe oxidative stress [34]. Another protein that illustrates an important influence on priming response is RecA, with the mutant showing a decreased priming response to H_2_O_2_. Overall, our data indicate that the priming by a low dose of H_2_O_2_ is multifactorial, with several OxyR-controlled proteins such as KatG, AhpCF or other factors related to DNA as RecA contributing to the priming response.

### Priming response enhances the survival of evolving populations

To find out whether the priming response described above has an influence on the rate of evolution of resistance to H_2_O_2_, we used an experimental evolution approach. We evolved bacterial cultures with a treatment protocol described in detail in the M&M section. Parallel populations were evolved where one group was periodically exposed to a sub-lethal concentration of H_2_O_2_, an activating signal that should protect the populations in comparison with naïve ones when later exposed to a higher dose. We continued daily intermittent exposures to H_2_O_2_ doubling both, priming and triggering concentrations until extinction occurred. We consider that a population is extinct when it shows no sign of growth during the next passage. This was also confirmed by periodical contamination checks after each passage. We observed that the extinction rate was faster for the naïve populations in comparison to primed populations (Log-rank test, *p*< 0.05, Fig 7). Naïve cells evolved resistance allowing them to resist up to 8 mM H_2_O_2_, whereas primed cells evolved resistance to survive up to 16 mM H_2_O_2_. These results show that priming increases the evolvability of pre-treated populations. We repeated the selection experiment including an additional control to exclude the possibility that primed populations evolved better because they received 10 % more H_2_O_2_ (priming plus triggering concentrations together). The result of this experiment showed a similar pattern to the previous one, naive populations went extinct first, including the one receiving ten percent more H_2_O_2._ Both naive populations groups were not significantly different (S3 Fig.).

**Fig 7.**
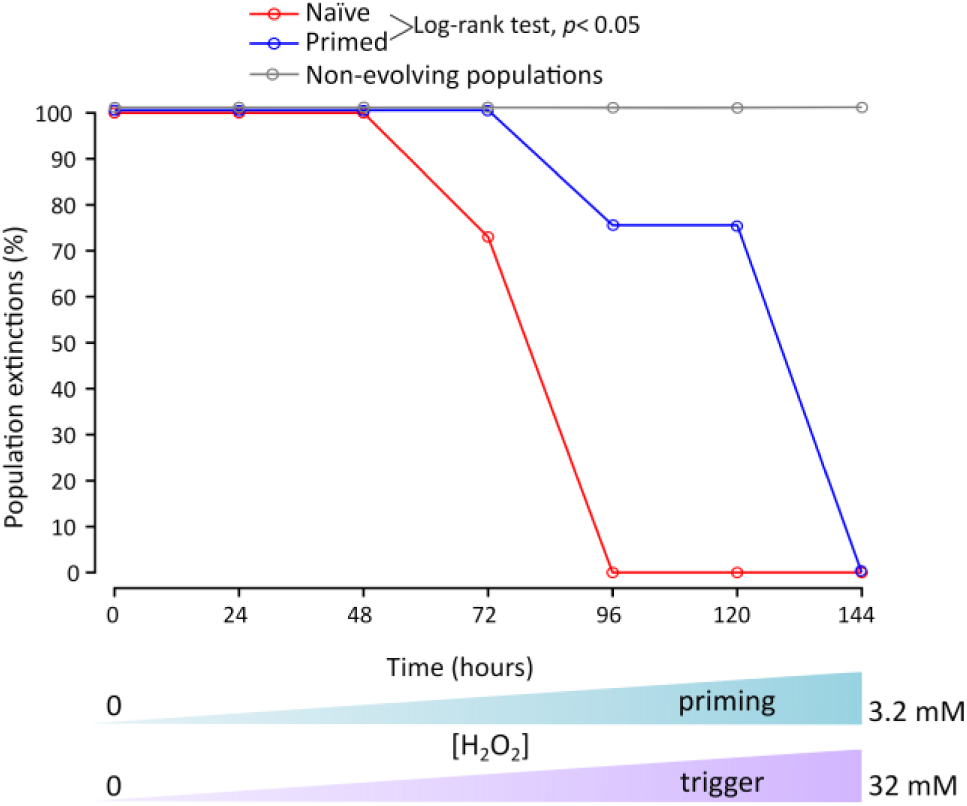
Proportion of population extinctions (20 populations per treatment) due to exposure to increasing concentrations of H_2_O_2_ during experimental evolution of 40 individual populations. Note that every passage was carried out every 24 hours although time in the x-axis is represented as continuous. Both, priming and triggering doses were increased twofold daily up to 32 mM, where total extinction occurred. The extinction was perceived by negative growth in the next passage and by the absence of growth In LB plates during contamination controls. Non-evolving population control (grey line, 20 populations) is also shown. Evolvability differs between the two treatments, naïve (red line) and primed (blue line) populations (Log-rank test, *p*< 0.01). Differences with non-evolved populations were not determined.

Further analysis of the resistant populations by whole-genome sequencing revealed that all populations harboured different sets of mutations. Surprisingly, we did not find any direct modification in the enzymatic scavenger systems, such as catalase or any other proteins related to peroxide protection, but we cannot exclude changes in expression level of these systems due to regulatory mutations acting *in trans*. There are hundreds of single nucleotide polymorphisms (SNPs) and other types of mutations that were unique to each population for both regimes (Fig 8). Many of these mutations probably represent neutral or non-lethal changes that populations accumulated during the exposure to H_2_O_2_. Here, H_2_O_2_ is not only a selective agent, but it also speeds up evolution by increasing mutagenesis. At the moment, we cannot be certain about the contribution of particular mutations to H_2_O_2_ resistance and they will be subject to detailed studies in the future. However, we studied two cases of the most frequent mutations in more detail as a proof of principle.

**Fig 8.**
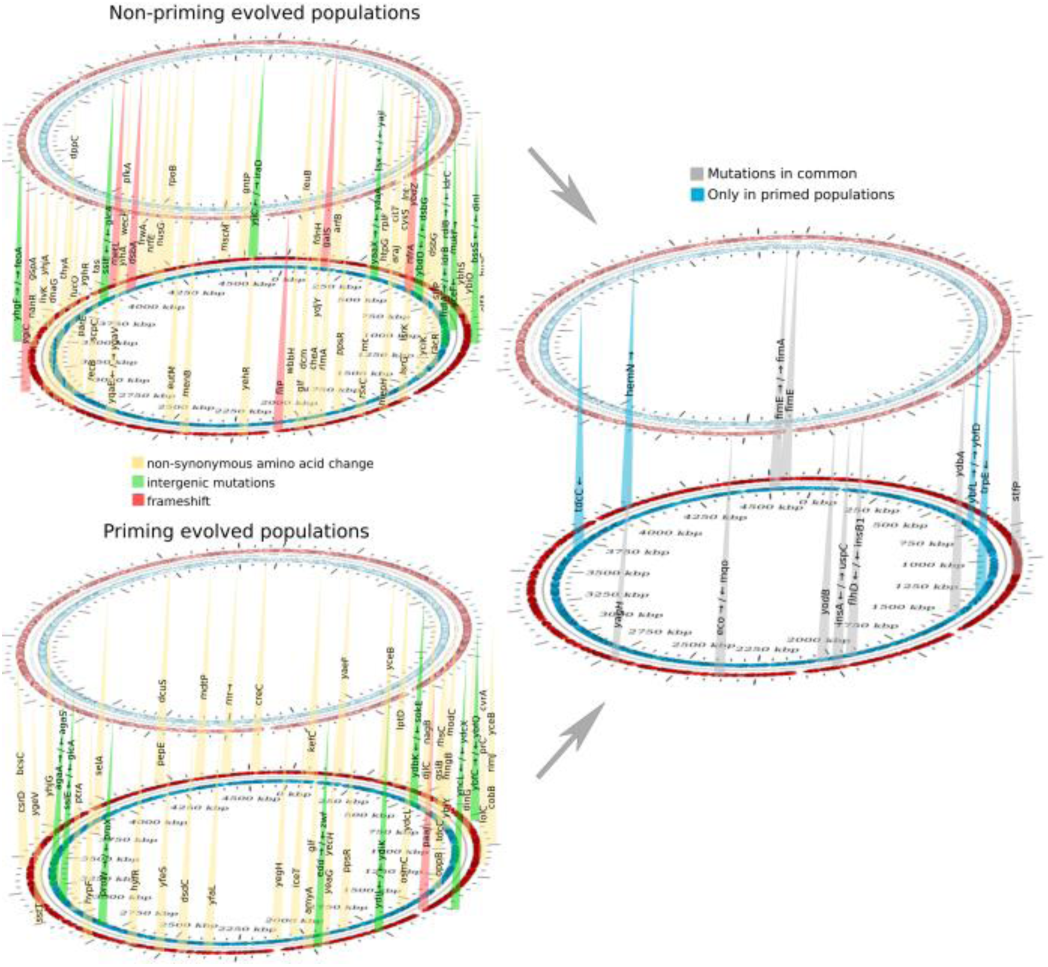
Chromosomal mapping of mutations in H_2_O_2_-evolved populations. The positions indicate the approximate locations of mutations in the *E. coli* MG1655 chromosome relative to the wild-type. Different types of mutations that were found in both evolving regimes are indicated in the left panel highlighted with different colours (non-synonymous amino acid changes in yellow, intergenic mutations in green and frame-shift in red). The right panel indicates mutations that were present for both regimes (grey) and those that were exclusive to the priming regime (blue). To see full reports of mutation for each population see supplementary material.

One first case is that of very frequent inactivating mutations in *fimE*. The promoter of the fimbrial subunit gene, *fimA* (the first gene of the operon *fimAICDFGH*) lies within a short segment of invertible DNA known as the *fim* switch (*fimS*), and the orientation of the switch in the chromosome determines whether *fimA* is transcribed or not. Inversion is catalysed by two site-specific recombinases, the FimB and FimE proteins. The FimB protein inverts the switch in either direction, while FimE inverts it predominantly to the off orientation. When FimB and FimE are co-expressed, FimE activity dominates and the switch turns to the off phase, wherease a fimE knockout mutation increases fimbriae production [35].

We selected one of the many *fimE* defective mutants (Δ1 bp, position 248 out of 597 nucleotides) for the next experiments. This type of mutant was only present in primed populations although there were some other intergenic mutations between *fimE*→*fimA*, both in primed and non-primed populations, pointing to the relevance of this type of mutation in survival under H_2_O_2_ stress. A series of experiments showed that *fimE* mutants attach to glass surfaces more efficiently than the wild-type strain, suggesting that production of fimbriae is active in the mutant. Furthermore, the *fimE* mutant showed decreased susceptibility to H_2_O_2_, with an MIC of 4 mM in compared to 1 mM in the parental strain. The transformation of the mutant strain with a plasmid overexpressing *fimE* (pCA24N-*fimE*, GFP minus) [36] reverted the attaching ability (Fig 9) of the mutant and also restored the original resistance to 1 mM H_2_O_2_. In early biofilm research it was shown that type I fimbriae (the product of the *fim* operon) are required for submerged biofilm formation. Type I pili (harbouring the mannose-specific adhesin, FimH) are required for initial surface attachment [37]. Fimbriae expression *per se* constitutes a signal transduction mechanism that affects several unrelated genes via the thiol-disulfide status of OxyR [38]. Fimbriae formation is accompanied by massive disulfide bridge formation [38] that could also contribute to titration of the exogenous H_2_O_2_, thereby limiting the intracellular damage.

**Fig 9.**
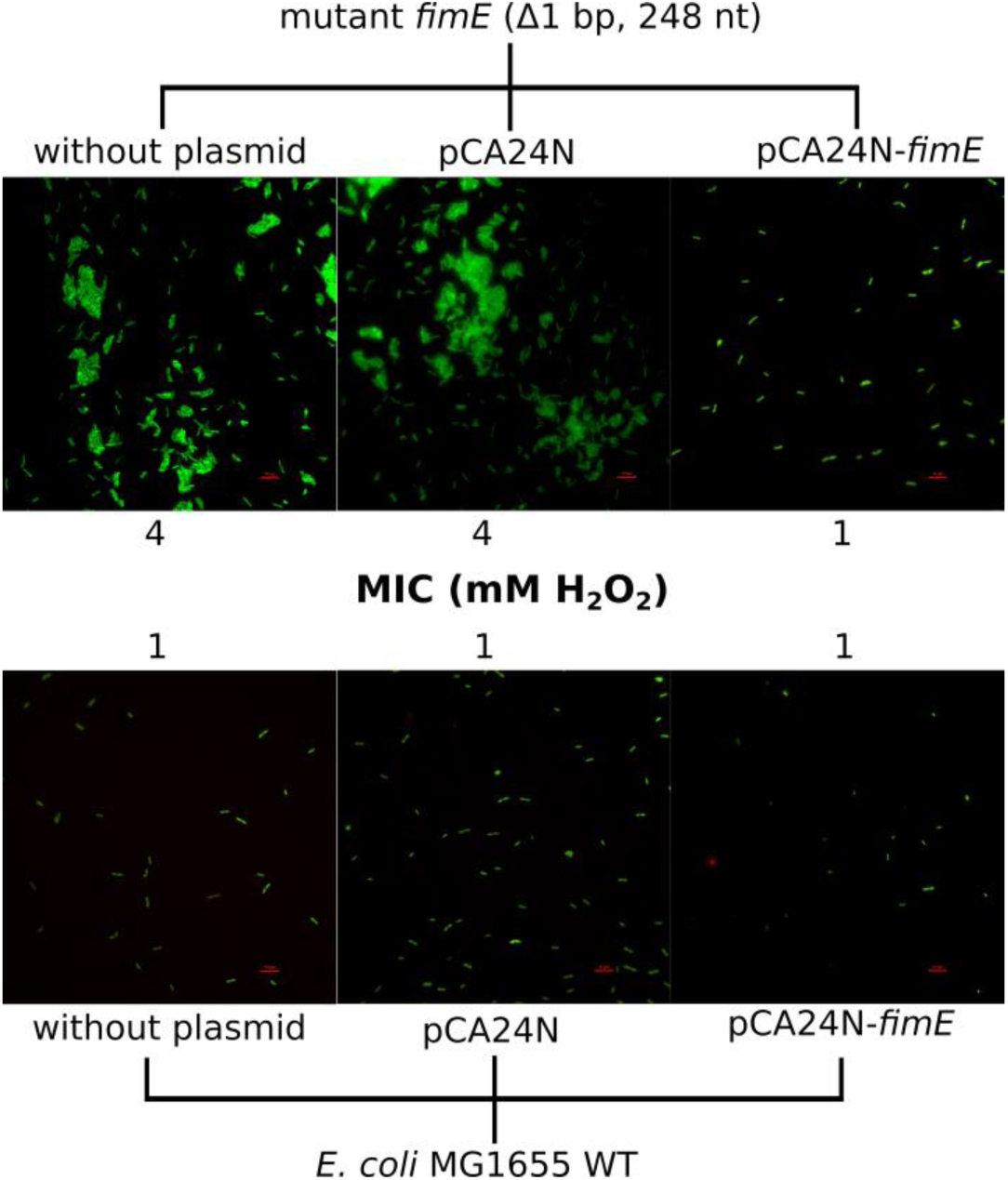
Representative fluorescent micrographs of attachment to a glass slide by *E. coli* MG1655 and its derivative mutant *fimE* (Δ1 bp, position 248 out of 597 nucleotides). The image was taken after vital staining (LIVE/DEAD BacLight Bacterial Viability Kit). Top panel shows greater attachment and reduced sensitivity to H_2_O_2_ MIC associated with *fimE* inactivation. Complementation restores both H_2_O_2_ sensitivity and the low-attachment phenotype of the wild-type strain (lower panel). In both panels control cells (transformed with the cloning vector pCA24N) can be also observed.

Following the inversion of the phase switch to ‘on state’ of fimbriae production by environmental signals, this element can remain phase-locked in the ‘on orientation’ due to integration of insertion sequence elements at various positions of the *fimE* gene [39]. Interestingly, *fim* operon expression allows *E. coli* to attach to abiotic surfaces, host tissues and to survive better inside macrophages protecting against the presence of extracellular antibacterial compounds [40,41]. Reactive oxygen species (ROS) are critical components of the antimicrobial repertoire of macrophages to kill bacteria [42].

A second case that we studied is the intergenic mutations between the genes *insB1* and *flhD (insB1*→*flhD)*, which occurred at high frequency in our evolution experiment. FlhD is coexpressed from an operon with FlhC and both proteins form the master transcriptional factor that regulates transcription of several flagellar and non-flagellar operons by binding to their promoter regions [43]. It is known that in *E. coli MG1655*, some insertion sequences such as insB1 can increase the motility of *E. coli* [44]. Our hypothesis here was that mutations in the intergenic region between *insB1 and flhD* contribute to abolishing the positive effect of *insB1* insertions on motility, which in turn increases resistance to H_2_O_2_.

We first analysed whether mutants from our evolution experiment showing these mutations showed decreased motility. Two independent mutants (positions 1978493 nt, Δ10bp and 1978504 nt, Δ1 bp) from two different populations showed decreased swimming and increased sedimentation when culture tubes were not shaken, as was the case in the evolution experiment setup. Both mutants also showed a MIC of 4 mM to H_2_O_2_, compared to 1mM in the control. The transformation with a plasmid overexpressing the operon *flhDC* (pVN15) restored both motility and H_2_O_2_ sensitivity (Fig 10). These mutations may protect against H_2_O_2_ through the mechanism of decreased motility, which results in clustering of the cells at the bottom of the culture, which may improve protection of cells located inside the clusters.

**Fig 10.**
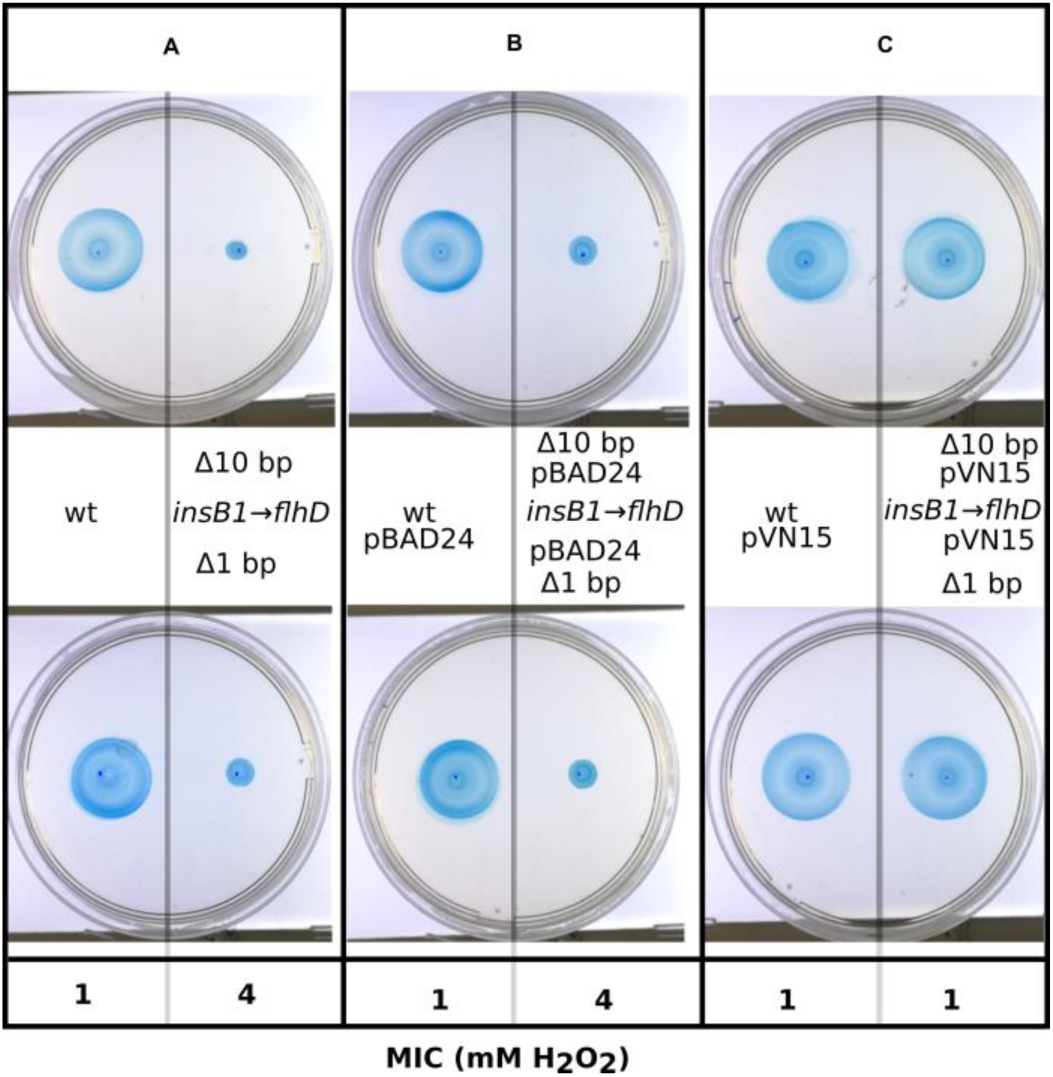
Motility test of *E. coli* MG1655 and its derivative mutants in the intergenic region between *insB1* and *flhD (insB1*→*flhD*, positions 1978493 nt, Δ10bp and 1978504 nt, Δ1 bp). Panel A shows the initial phenotypes of *insB1*→*flhD mutants*, low motility and resistance to 4 mM H_2_O_2_ while the parental strain is motile with a MIC of 4 mM H_2_O_2_. A complementation experiment showed identical phenotypes when the strains were transformed with the cloning vector (panel B) and recovery of motility and original resistance to 1 mM H_2_O_2_ when both mutants are transformed with the plasmid overexpressing the operon *flhDC* [80] (panel C).

A second possibility is that the decrease of motility itself decreases the basal level of H_2_O_2_ due to lower metabolic demand. Flagellar motility enables bacteria to escape from detrimental conditions and to reach more favourable environments [44]. However, flagella impose an important energetic burden on bacterial metabolism due to the number of proteins involved in the machinery as well as the associated energy expenditure associated with motility. For instance, one interesting study showed improved tolerance to oxidative stress in *Pseudomonas putida* as reflected by an increased NADPH/NADP(+) ratio, concluding that flagellar motility represents the archetypal trade-off involved in acquiring environmental advantages at the cost of a considerable metabolic burden [45]. In our condition of oxidative stress, the flagellate phenotype makes the cells more susceptible to H_2_O_2_. These results raise an interesting question in regard to the motile vs non-motile strategy in bacteria: does flagellar activity bring diminishing returns by creating sensitivity to oxidative stress? The decreased expression in the flagellar gene hierarchy also affects *pdeH*, a class III gene in this hierarchy, which encodes the major c-di-GMP degrading enzyme in *E. coli*. The result is an increase in c-di-GMP levels, which promotes the production of curli fibers, which are a major component of the extracellular biofilm matrix. The strain MG1655 does not produce cellulose; strains that do so would have also increased cellulose production. Curli fibers and cellulose production are co-regulated by CsgD, which is itself under positive c-di-GMP control. Thus, the reduction on FlhDC level promotes biofilm formation, which contributes to multiple stress resistance, including resistance against H_2_O_2_ [46]. We can speculate about the role of some other mutations. For example, a set of changes are located in genes coding for iron-binding proteins or related to iron transport such as *iceT, feoA, yaaX*/*yaaA* or *rsxC*. The control of intracellular iron is crucial to decrease the adverse effects of Fenton chemistry [7]. There were also mutations that were common to both types of population, evolved under priming and non-priming conditions (Fig 8). The most frequent mutations were *yodB* (a cytochrome b561 homologue), intergenic mutations between *insA* and *uspC* (universal stress protein C). Another frequent mutation was in the gene *yagH*, belonging to the CP4-6 prophage. CP4-6 is a cryptic prophage in *E. coli* that could play a role in bacterial survival under adverse environmental conditions [47].

### Priming alters the mutational spectrum and H_2_O_2_-induced mutagenesis in evolving populations

To assess if the evolved populations have a similar mutational spectrum, we analysed the total pool of mutations segregated by the treatments. We used the Monte Carlo hypergeometric test implemented by iMARS [48] to assess the overall differences between each mutational spectrum. Both groups, evolved under priming and non-priming conditions, differed from each other significantly (*p*=0.00021). ROS induces a particular type of mutations, with a characteristic signature in the DNA. The guanine base in genomic DNA is highly susceptible to oxidative stress due to its low oxidation potential. Therefore, G·C→T·A and G·C→C·G transversion mutations frequently occur under oxidative conditions [49,50]. Thus, we investigated the proportion of C→A and C→G substitutions between the two types of evolving regimes, but we did not find significant differences (*p*=0.056 and *p*=0.11 respectively, two-tailed Fisher’s exact-test, Fig 11).

**Fig 11.**
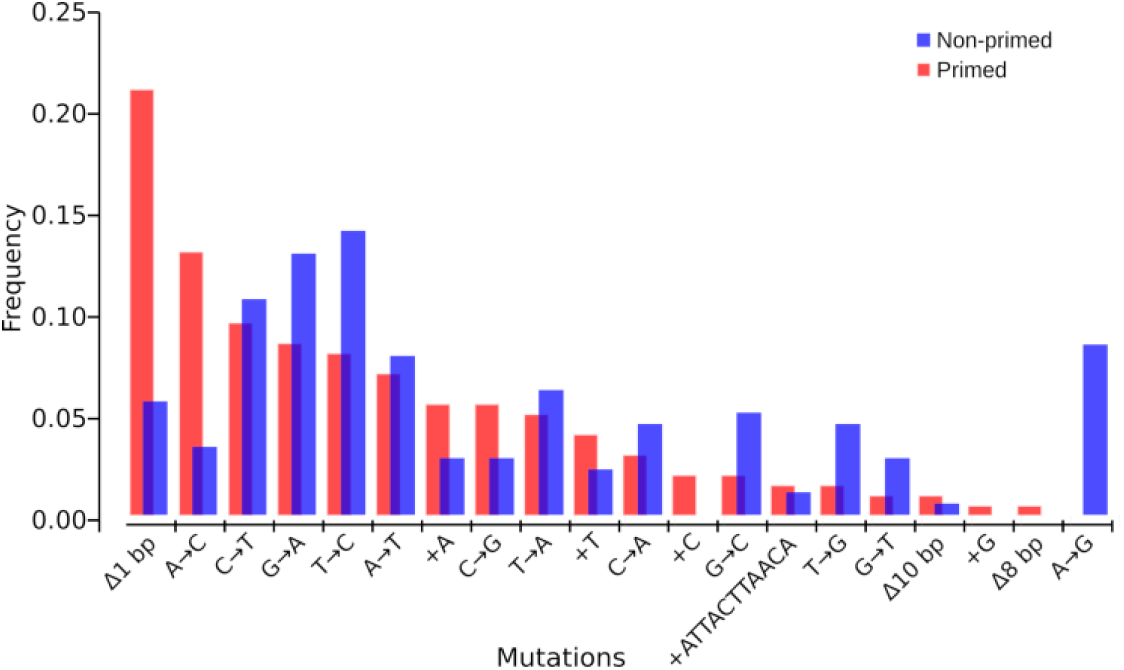
Mutational spectra. Frequencies of different types of mutations among evolved populations of both primed and non-primed conditions. The number of mutations (nucleotide substitutions and indels) is plotted against respective nucleotide positions within the gene fragment. In this analysis, the mutations were taken by type regardless of the targets by generating two frequency datasets to compare frequency by type of mutation of evolved populations under priming and non-priming regimes. The mutational spectra are significantly different between the two conditions according to the spectrum analysis software iMARS [46].

A possible explanation for these results is that the pre-activation of antioxidant defences helps to decrease the number of non-adaptive mutations and lethality by decreasing the extent to which the Fenton reaction occurs and the amount of released hydroxyl radicals. We observed an increase in the frequency of substitutions G→A, T→C, G→C, T→G and A→G in naïve populations compared to primed populations, indicating that the priming response greatly buffers DNA-damage. Primed populations showed an increased frequency of frameshift mutations by deletion of 1 bp, and additions of C or G that were only present in primed populations that also showed a higher number of A→C substitutions.

We hypothesised that the priming response would decrease DNA damage and hence also the rate of lethal mutations. To assess this possibility, we tested if priming can decrease the mutant frequency and hence lethality due to mutagenesis. Applying the conditions of the previous experiments, we determined the mutation frequency with and without H_2_O_2_ treatment. We found that the priming response decreased H_2_O_2_-induced mutation frequency close to one order of magnitude compared to naïve cells (*p*<0.001 primed versus naïve, Welch’s test, S5 Fig). In addition, there is a very small (1.87-fold change) but significant difference between primed bacteria and the basal *E. coli* mutagenesis (*p*=0.034 primed versus base level, Welch’s test), indicating that priming suppresses most of the mutagenesis caused by H_2_O_2_. In our experiment, naïve and primed populations showed different death rates (Fig 1). We also showed previously that different inoculum sizes in similar conditions to our current experiment do not influence mutagenesis [51]. We propose that one of the most important consequences of the priming response to H_2_O_2_ is a drastic decrease in lethal mutagenesis. Although H_2_O_2_ damages most of the cellular components [52], DNA damage is likely the major contributor to lethality.

In principle, an increase in mutation rate increases the evolvability of asexual populations [53–55]. How is it possible that naïve populations show lower evolvability compared to primed populations despite a higher mutation rate? A possible explanation is that evolvability can be influenced by the population size and the mutation supply. Even with increased mutagenesis, if population size drastically decreases, the final number of mutants can be smaller when survival is improved. We also found that the mutational spectra of our evolving populations are different. Mutational spectra are a qualitative property of mutation rate that could enhance or hinder the access to beneficial mutations [56]. It is possible that the observed changes in mutational spectra between primed and naïve evolving populations could play a role in balancing the ratio of deleterious/beneficial mutations, although demonstrating this will be the subject of future research.

## Conclusions

Priming by H_2_O_2_ in exponentially growing bacteria is quantitatively driven by genes that are mostly under OxyR control, with other genes such as *recA* also contributing to survival. The memory of the priming response can last up to four generations upon first exposure to H_2_O_2_. We also showed high stability of H_2_O_2_-detoxifying proteins, which play a significant role in resistance.

How general priming is and to which other stressors it applies remains to be seen. Our findings are also contributing to an understanding of ROS-mediated interactions of hosts with pathogens and the microbiota.

Our finding that priming boosts evolvability of bacterial populations by enhancing survival under oxidative stress, provides evidence for the phenotype as a target of selection during evolutionary processes and supplies a validation for the ‘plasticity-first’ hypothesis [4]. If such selection happens in the more complex environment of a host remains to be studied. Bacteria, however, are certainly frequently exposed to fluctuating concentration of ROS, concentrations that often will be sublethal.

The type and number of mutations indicate that scavenger systems against oxidative stress are optimally evolved since no mutations directly affecting these systems were found under H_2_O_2_ stress selective pressure. Our results suggest that the ubiquitous occurrence of H_2_O_2_ has an impact on bacterial lifestyle and the evolution and regulation of flagella motility. Moreover, mutations that result in bacterial clustering or increase bacterial density can contribute to protection against H_2_O_2_ as shown by mutations of the flagella regulator or in the *fim* operon. It seems possible that H_2_O_2_ thereby stimulates the early stages of biofilm formation, thus providing additional protection against ROS.

## III. Material and methods

### Bacteria and growth conditions

*E. coli* MG1655 was used as bacterial model for all experiments with H_2_O_2_. For genetic manipulation, *Escherichia coli* strain DH5α was used and routinely cultured in Lysogeny Broth (LB medium), supplemented with antibiotics when appropriate. All bacterial strains were cultured in Lysogeny Broth Lenox (Carl Roth, Germany). For all experiments with H_2_O_2_ the LB was freshly prepared and kept in the dark until use.

### Construction and verification of deletion mutants

All mutants used in this work (S9 Table) were generated in *E. coli* K-12 strain MG1655 following a modified methodology described elsewhere [57]. Briefly, transformants carrying the red recombinase helper plasmid, pKD46, were grown in 5-ml SOB medium with ampicillin (100 µg/ml) and L-arabinose at 30°C to an OD_600_ of ∼0.5 and then made electrocompetent. PCR products with homology regions were generated using specific primers (S10 Table) to amplify the region of interest from the corresponding mutants of the Keio collection [58]. The PCR-generated fragments were purified (MinElute PCR Purification Kit, Qiagen). Competent cells in 50 µl aliquots were electroporated with 100 ng of PCR product. Cells were added immediately to 0.9 ml of SOC, incubated 1 h at 37°C, and then 100 µl aliquots spread onto LB agar with kanamycin (30 µg/ml). The mutants were verified by PCR and the antibiotic resistance cassette was removed using the plasmid pCP20. The correct inactivation of genes was verified by PCR. To construct double mutants, single mutants obtained in MG1655 were transduced using the P1vir phage procedure as previously described [59].

### Hydrogen peroxide susceptibility testing

We determined the minimal inhibitory concentration for H_2_O_2_ by broth microdilution method with some modifications. We used LB medium instead of Mueller-Hinton Broth and we used approximately 10^7^ bacteria instead of 10^5^, a bacterial density that corresponds to the subsequent experiments. When working with H_2_O_2_, we carried out all experiments with freshly-made LB to avoid the accumulation of chemically formed H_2_O_2_ by the joint action of light and Flavin [60] present in the medium during storage. Time-kill curves to H_2_O_2_ were determined by exposing exponential phase bacteria at a density of ∼ 2×10^7^ CFU/ml to different concentrations and times, taking as a reference the modified MIC value.

### Priming experiments with H_2_O_2_

Starting from 2×10^7^ CFU/ml, *E. coli* was exposed (stimulus) to 0.1 mM H_2_O_2_ during 30 minutes at 37°C with shaking. The H_2_O_2_ was removed by centrifugation at 4 000 x g for 10 minutes and cells were allowed to recover for 90 minutes, keeping the cell density constant by removing the required volume and replacing it with the appropriate amount of fresh pre-warmed-LB (37°C) every 30 minutes. The trigger (1 mM H_2_O_2_) was added 90 minutes after removal of the stimulus. The challenge lasted for 30 minutes. At this point, 4 μg/ml catalase (Sigma Aldrich, Germany) was added to each tube and cultures were diluted and plated to determine cell viability. Non-treated cells were used as control. Each group consisted of five cultures.

### Determination of the priming cost

Bacterial growth curves were measured in flat-bottom 96-well micro-plates (Nunc, Denmark). 40 independent colonies from a LB agar plate were inoculated in a 96 multi-well plate containing 200 μl/well of LB and incubated overnight at 37 °C in a humid chamber to prevent evaporation. In a new plate, each well was filled with 100 μl of fresh-made LB and inoculated per duplicate with 1 μl from the overnight plate (80 wells in total). The remaining 16 wells were used as medium contamination control. The plate was incubated for 2.5 hours to reach an OD_600_ of 0.4. Then, 90 μl of medium per well were removed and replaced with 80 μl of fresh LB using a multichannel pipette. Ten additional microliters containing 1 mM H_2_O_2_ were added to every 40 wells and mixed immediately. Only LB was added to the remaining 40 control wells. Each independent colony was represented in both experimental groups. The plate was placed into a microplate reader Synergy H1 (Biotek, Germany) and kinetic readings (OD_600_) were carried out every 20 minutes after short shaking of 5 seconds. The cost of priming by H_2_O_2_ (0.1 mM) was estimated from the parameters of the growth curves. All model parameters —carrying capacity, initial population size, growth rate, doubling time and the empirical area under the curve— were calculated using Growthcurver R package [61]. Each parameter for *E. coli* MG1655 growing with or without 0.1 mM H_2_O_2_ (priming concentration), was compared using Student’s t-test.

### Hydrogen peroxide colourimetric quantitation in *E. coli* supernatant media

Pierce Quantitative Peroxide Assay Kit (Thermo Scientific, Germany) was used according to manufacturer’s instructions. Briefly, *E. coli* cells were grown in fresh LB (Lennox) medium to OD_595_ 0.5. The cultures were diluted 10 times in 1 ml of fresh LB containing 0.1 mM H_2_O_2_ (final concentration, primed) or LB alone (naïve). The tubes were incubated with moderate shaking for 30 minutes. Then, the supernatant was removed by centrifugation at 10000 x g during 1 minute and aspiration. The pellets were washed once with LB and resuspended in 1 ml of LB containing 1 mM H_2_O_2_. At time points 0, 15 and 30 minute 100 μl of supernatant were removed to determine the H_2_O_2_ concentration. Finally, 20 μl microliters of medium supernatant per tube were diluted 10 times and mixed in a 96-well microplate with 200 μl of working solution (prepared according to the manufacturer’s instructions). The mix was incubated at room temperature for 15 minutes in a humid chamber. OD_600_ was measured using a Synergy H1 plate reader (Biotek, Germany) and a standard curve of H_2_O_2_ was prepared as indicated in the protocol. The blank value (working without H_2_O_2_) was automatically subtracted from all sample measurements. Three independent cultures were used per group and non-treated cultures were used as an additional control. Means and standard deviations were calculated and compared for each time-point by a Welch’s test.

### Memory of the priming response

After applying a stimulus (0.1 mM H_2_O_2_) for 30 minutes, the cells were allowed to recover for 30, 60, 90, 120 and 150 minutes before the addition of the trigger concentration (1 mM H_2_O_2_). During the experiment, the OD_600_ of all cultures were kept around 0.05 by adding suitable volume and removal of equivalent quantity to maintain the population in the exponential phase and the same population size. Bacteria remained exposed to the trigger for 30 minutes before starting the dilution in 1 ml of LB containing 4 µg of catalase. Appropriate dilutions were made and plated onto LB agar to determine the survival rate.

### Global proteomics by LC-mass spectrometry

*E. coli* strain MG1655 was grown at 37°C in LB medium to an OD_600_ of 0.5. The cultures were diluted 10 times in fresh LB. H_2_O_2_ was added to tubes to final concentrations of 0.1 and 1 mM. Non-treated samples were used as control. All tubes were incubated for 5 minutes with shaking at 37°C. Remaining H_2_O_2_ was removed by adding 4 µg/ml of catalase first and by centrifugation at 10 000 x g during 2 minutes. After removal of the supernatant, the equivalent amount of fresh LB was added. For the memory decline experiment, the procedure was identical, except that only 0.1 mM H_2_O_2_ was used and the samples were treated for 30 minutes. After removal of the treatment, samples were taken after 30, 60, 90, 120 and 150 minutes. Each experimental condition consisted of six replicates. One millilitre per sample was pelleted by centrifugation at 10 000 x g for 2 minutes. The cells were resuspended in 50 µl of TE (10 mM Tris-HCl pH 8.0, 1 mM EDTA) containing chicken lysozyme (0.1 mg/ml, Sigma Aldrich, Germany) and incubated at room temperature for 5 minutes with occasional swirling. A volume of 250 µl of denaturation buffer (6M urea/2 M thiourea in 10 mM HEPES pH 8.0) was added into each sample and 25 µl (which approximately corresponds to 50 µg of protein) of the resulting lysate were used for in-solution protein digestion as described previously [62]. Briefly, proteins, re-suspended in denaturation buffer, were reduced by the addition of 1 µl of 10 mM DTT dissolved in 50 mM ammonium bicarbonate (ABC) and incubated for 30 minutes, followed by 20-minute alkylation reaction with 1 µl of 55 mM iodoacetamide. As a first digestion step, Lysyl endopeptidase (LysC, Wako, Japan) resuspended in 50 mM ABC was added to each tube in a ratio of 1 µg per 50 µg of total proteins and incubated for 3 hours. After pre-digestion with LysC, protein samples were diluted four times with 50 mM ABC and subjected to overnight trypsin digestion using 1 µg/reaction of sequencing grade modified trypsin (Promega, USA), also diluted before use in 50 mM ABC. All in-solution protein digestion steps were performed at room temperature. After the addition of iodoacetamide, the samples were protected from the light until the digestion was stopped by acidification adding 5% acetonitrile and 0.3% trifluoroacetic acid (final concentrations). The samples were micro-purified and concentrated using the Stage-tip protocol described elsewhere [62], and the eluates were vacuum-dried. Re-dissolved samples were loaded on a ReprosilPur C18 reverse phase column and peptides were analysed using a nano-HPLC Dionex Ultimate 3000 system (Thermo Scientific, Germany) coupled to an Orbitrap Velos mass spectrometer (Thermo Scientific, Germany). MS and MS/MS data from each LC/MS run were analysed with MaxQuant software [63]. Identification of proteins was performed using the MaxQuant implemented Andromeda peptide search engine and statistical analysis was carried out using the software Perseus [64].

### Prediction of *in vivo* protein half-life and stability index

The half-life estimation is a prediction of the time that it takes for half of the amount of protein in a cell to disappear after its synthesis. It relies on the “N-end rule” (for a review see [65–67]). The instability index provides an estimate of the stability of a protein in a test tube. Based on experimental data [68], making possible to compute an instability index using the amino-acid sequence. For these predictions, we used the online software ProtParam [69]. When available, N-end sequence was corrected to the real *in vivo* sequence due to methionine excision [70].

### Quantification of gene expression

To quantify the expression of H_2_O_2_ stress related genes in *E. coli*, bacteria treated with 0.1 mM H_2_O_2_ were compared to an untreated control. Preparation of each of the six treatments and the control samples were carried out by two operators three times on three consecutive days, thereby comprising three biological replicates for each of the six experimental and one control groups. Overnight (ON) cultures were diluted 1:100 in fresh LB, subdivided into six 10 ml aliquots and incubated in 50 ml-Falcon tubes at 37°C for 2.5 hours at 220 R.P.M. When the OD_600_ reached 0.4-0.5, the cultures were diluted 10 times with LB and H_2_O_2_ was added to the final concentrations of 0.1 mM. These bacterial cultures were incubated for 30 minutes at 37°C with shaking. Catalase was added to each tube to a final concentration of 4 µg/ml. Ten millilitres of treated and control cultures were collected, immediately centrifuged at 10 000 x g for 2 minutes and the supernatant was removed. The bacterial pellets were resuspended in 1 ml LB and mixed with 1 ml of RNAprotect Bacteria Reagent (Qiagen, Germany), incubated during 2 minutes and centrifuged at 10 000 x g for 2 minutes at room temperature. The supernatant was discarded and the bacterial pellet was immediately frozen and stored at −80°C until RNA extraction.

RNA was isolated using RNeasy kit (Qiagen, Germany) according to the manufacturer’s instructions and eluted in 50 μl of RNase-free water. The nucleic acid yield and purity were determined by measuring the optical density at A260/280 using Nanodrop spectrophotometer (Thermo Scientific). RNA samples were treated with TURBO DNase (Life Technologies, Germany). Briefly, 10 μg of RNA were used in a total volume of 500 μl containing 20 units of TURBO DNase, incubated for 30 minutes at 37°C, immediately followed by RNeasy (Qiagen, Germany) clean-up and elution in 30 μl of RNase-free water. Following DNase treatment, RNA integrity was assessed using Agilent RNA 6000 Nano kit and 2100 Bioanalyzer instrument (both Agilent Technologies, USA). All samples had RIN values above 8.

For cDNA synthesis, total RNA (250 ng per reaction) and random primers were used for cDNA synthesis using High-Capacity cDNA Reverse Transcription Kit with RNase Inhibitor (Applied Biosystems, Germany). Initially, to ensure linear conversion of the transcripts, the dynamic range of reverse transcription reaction was tested by performing a standard curve with cDNA, synthesised using various input amounts of pooled RNA. To obtain a sufficient amount of cDNA, several batches of 20 μl RT reactions were pooled, diluted 50-fold with RNase-free water and stored in single-use aliquots at −80°C until further use. All 21 samples were tested for presence of contaminating genomic DNA by running the *mdoG* assay with cDNA and the respective no reverse transcription (-RT) controls. There was no amplification in the majority of –RT controls. In –RT samples with detectable amplification, difference in Ct values when compared with +RT varied between the samples, but was no less than 10 cycles for all of them with the lowest Ct values in –RT control samples ≥ 30.

For primer design, *Escherichia coli* strain K-12 MG1655 complete genome (accession U00096) sequence was downloaded from NCBI database (www.ncbi.nlm.nih.gov) and used as reference. Target sequence accession number, and primer sequences for each assay can be found in S11 Table. Primers were designed using Primer Express software (Applied Biosystems, Germany) and optimised for annealing temperature of 60°C. Each primer pair and amplicon were checked for secondary structure formation using Oligo Tool (Integrated DNA technologies, USA, http://eu.idtdna.com/analyzer/Applications/OligoAnalyzer/).

Quantitative Real-time PCR reactions were prepared manually in a total volume of 10 μl by mixing 5 μl 2x KAPA SYBR FAST ABI Prism master mix (KAPA Biosystems, Germany), 0.2 μl forward and reverse primer mix (10 μM each primer), 2.8 μl RNase-free water and 2 μl cDNA in MicroAmp Fast Optical 48-wells reaction plates (Applied Biosystems, Germany). PCR reactions were set up in the Fast mode using StepOne thermocycler (Applied Biosystems, Germany) with the following cycling conditions: 95°C 3’, 40X (95°C 3’’, 60°C 20’’), melting curve analysis. Each assay was run in duplicate. No template controls (NTC) were included each time. Presence of a single specific product was verified by running a melt curve analysis followed by visualisation of the qPCR product in a 2% agarose gel stained with SYBR Safe DNA Gel stain (Life Technologies). Additionally, each PCR assay was tested for reaction efficiency as follows: equimolar amounts of cDNA from all 21 samples were pooled together and used for the preparation of the standard curve by serial dilution (1:3) over five dilution points and run in triplicate. Expression of target genes was normalised to the expression levels of three reference genes (*arcA, mdoG* and *tus*), selected based on the assessment of the expression stability across all experimental conditions using BestKeeper software [71]. Reaction efficiency information inferred from the standard curve data was used to correct for differences in amplification efficiencies in the REST 2009 software [72]. Default settings (2000 iterations) were used for randomisation and bootstrapping analysis to test significance of gene expression. Expression values with *p*-values ≤0.05 were assigned as differentially expressed. We followed the Minimum Information for Publication of Quantitative Real-Time PCR experiments (MIQE) guidelines-compliant check-list [73].

For data analysis, qPCR Amplification curves were first visually examined in the StepOne software (Applied Biosystems, Germany). No baseline and threshold line adjustments were necessary. Ct values of the technical replicates were averaged and used for relative gene expression analysis in the REST 2009 software (Qiagen, Germany) [72]. Expression of target genes was normalised to the expression levels of three reference genes (*arcA, mdoG* and *tus*), selected based on the assessment of the expression stability across all experimental conditions using BestKeeper software [71]. Reaction efficiency information inferred from the standard curve data was used to correct for differences in amplification efficiencies in the REST 2009 software. Default settings (2000 iterations) were used for randomisation and bootstrapping analysis to test significance of gene expression. Expression values with *p*-values ≤0.05 were assigned as differentially expressed. We followed the Minimum Information for Publication of Quantitative Real-Time PCR experiments (MIQE) guidelines-compliant check-list [73].

### Transcriptome sequencing

The transcriptome sequencing was carried out on samples treated with 0.1 mM H_2_O_2_ during the experiment of memory decline. The time point used corresponded to 120 minutes after removal of the treatment. Bacterial cells that were kept in low density (0.05 OD_600_) were concentrated 10 times and the pellets were resuspended in 1 ml of lysis buffer. Both, small RNA fraction and the large one were isolated using the microRNA & small RNA Isolation kit (Thermo Scientific, Germany). Traces of genomic DNA were removed from 10 µg of RNA by digestion in a total volume of 500 μl containing 20 units of TURBO DNase, incubated for 30 minutes at 37°C, immediately followed by RNeasy (Qiagen, Germany) clean-up and elution in 30 μl of RNase-free water. Following DNase treatment, RNA integrity was assessed using Agilent RNA 6000 Nano kit and 2100 Bioanalyzer instrument (both Agilent Technologies, USA). Both fractions were depleted from ribosome RNA using the Ribo-Zero Depletion Kit for Gram-negative bacteria (Illumina, USA). Libraries were prepared using a TruSeq Stranded Total RNA library preparation kit (Illumina, USA) and were sequenced on a MiSeq platform. All sequences are available from the NCBI SRA under BioProject accession PRJNA485867.

### Evolution experiment

Before the beginning of the experiment, five independent clones of *E. coli* MG1655 were pre-adapted to the experimental conditions such as culture medium, temperature and oxygen level. The pre-adaptation was carried out by diluting an overnight culture of each clone 1:1000 followed by incubation at 37°C in the plate reader with the same shaking conditions used during the rest of the experiment in 0.2 ml in fresh-made LB. The bacteria were cultured by serial passage every 24 hours using a bottleneck of 1% (1/100 dilution) to allow for approximately fifty generations during 10 days. Contamination checks by plating out on LB and cryopreservation of culture aliquots at −80°C in LB containing 15% glycerol solution were carried out periodically. All lines showed similar fitness and five independent colonies (one per line) were selected for the evolution experiment. The experiment was performed at 37°C with periodical shaking in a microplate reader (Synergy H1, Biotek, Germany). We used flat-bottom polystyrene 96-well plates which lids had anti-evaporation rings (Greiner Bio-One, Germany). A final volume of 200 µl per well was used and we founded 60 independent populations: 20 populations that we evolved under priming conditions, 20 populations under non-priming conditions and 20 non-evolving populations that were only serially passed. Growth curves were generated by taking measurements of OD_600_ every 20 minutes (preceded by a brief shaking of 5 seconds). Every day, the experiment consisted of two stages. Every morning a V-bottom plate containing 190 µl of LB was inoculated with 2 µl of overnight culture and incubated for 2 hours to reach an OD_600_ between 0.2 to 0.6. At this point, 3 µl of LB containing the corresponding priming concentration of H_2_O_2_ (for populations to be primed) or only LB (control populations) were added to each well. The plate was then incubated for 30 minutes, centrifuged and the supernatant removed using a Costar 8-channel vacuum aspirator (purchased from Sigma Aldrich, Germany) equipped with disposable tips. The pellets were resuspended in 200 µl of fresh LB and 190 µl were transferred to a new flat-bottom plate. Cells were allowed to recover for 30 minutes at 37 °C and challenged with a ten-fold higher concentration of the priming one. The dish was placed in the plate reader and the growth was followed as described above during 20 hours. The serial passage started at 50 µM of H_2_O_2_. Next day, the procedure was identical for priming and triggering concentrations that were doubled every 24 hours to reach a final challenging concentration (trigger) of 32 mM H_2_O_2_ where all bacterial population went extinct. Before each passage, 20 µl per population were added to 180 µl of sterile 0.9% NaCl and then serially diluted, plated and inspected for contamination and population extinction.

### DNA isolation

Genomic DNA samples for whole genome sequencing were isolated using in house method based on fast phenol:chloroform extraction, removal of RNA and ethanol precipitation. The DNA quantity and quality were estimated by measuring the optical density at A260/280 using a Nanodrop 2000 (Thermo Scientific, Germany) and agarose gel electrophoresis.

### Genome re-sequencing

We sequenced the total genomic DNA from sixty populations from the evolution experiment, all from the final passage before extinction (20 evolved from the priming regime, 20 from the non-priming regime and 20 control strains). TruSeq DNA PCR-free libraries were constructed according to the manufacturer’s instructions and sequenced for 600 cycles using a MiSeq at the Berlin Center for Genomics in Biodiversity Research. Sequence data are available from the NCBI SRA under BioProject accession PRJNA485867. The haploid variant calling pipeline snippy [74] was used to identify mutations in the selection lines. Snippy uses bwa [75] to align reads to the reference genome and identifies variants in the resulting alignments using FreeBayes [76]. All variants were independently verified using a second computational pipeline, breseq [77].

### Determination of H_2_O_2_-induced mutagenesis

This procedure was carried out following similar protocols to previous studies with some modifications [78,79]. Five independent cultures (5 ml each one) of *E. coli* MG1655 were grown in fresh LB medium to an OD_600_ of ∼0.2. Then, each culture was diluted 10 times (volume was increased to 50 ml) with LB and divided into two separate sets of 25 ml per culture each one. One set, consisting of five cultures of 25 ml was treated with 0.1 mM H_2_O_2_ (primed) while the second set of culture remained untreated (naïve). After 30 minutes, both cultures were treated with 1 mM H_2_O_2_ during another 30 minutes. Then, H_2_O_2_ was removed, first, by quickly adding 4 μl/ml of catalase and, second, by collecting the cells by centrifugation and washing them with one ml of fresh LB. Another set of tubes received no treatment and were used to determine the basal frequency of mutants. For determination of H_2_O_2_-induced mutation frequency, a 4 ml tube of fresh LB was inoculated with 1 ml of washed bacteria from each culture. The cultures were allowed to recover overnight for 16 h at 37°C. Serial dilutions of each culture were plated next day onto LB plates containing 100 µg/ml of rifampicin or without antibiotic to estimate the viability. The three groups were compared among them using a Welch’s test.

### Attachment assays

Attachment assays of an *E. coli* mutant in *fimE* and control strains to microscope glass slides were investigated. The experiment consisted of dipping sterile microscope slides into 50 ml Falcon tubes containing 10 ml of mid-exponential phase cultures of *E. coli* (0.5 OD_600_), five culture for each group. The slides were incubated during 1 hour. In the last 10 minutes of the incubation, 10 μl of the LIVE/DEAD^®^ BacLight Bacterial Viability Kit (Thermo Scientific, Germany) was added to each culture following the instructions of the manufacturer. The slides were mounted with a glass coverslip and fluorescent images were taken of several fields with a simultaneous acquisition in red and green fluorescent channels with a Nikon Ti-2 inverted microscope (Nikon, Japan). Cells were observed with the 100× objective and controlled by Nis Element AR software.

### Bacterial swimming assay

Swimming motility of mutants in the upstream region of *fldhC* and their complementation analysis were carried out on swimming plates containing swimming medium (0.5% bacto-tryptone and 0.3% agar, both from Sigma Aldrich, Germany). Briefly, 1 ml of overnight cultures per strain was centrifuged and resuspended in their supernatant adjusting their final OD_600_ to 4. From these OD-adjusted cultures, 2 μl from each one were inoculated into the swimming plates in pairs and bacteria were allowed to grow and swim for 4 h at 37°C. For complementation we used the plasmid pVN15 (pBAD24 carrying *flhDC* operon genes) and the vector pBAD24 was used as control [80]. For contrast purposes, another 5 ml of swimming medium containing 0.25 mM isopropyl-β-D-thiogalactopyranoside (IPTG) and 4 µg/ml 5-bromo-4-chloro-3-indolyl β-D-galactopyranoside (X-gal) were added carefully covering the entire surface of each plate. After the top layer solidification, the plates were incubated for 15 minutes at 37°C and 1 hour at 4°C before taking photographs. The combination of IPTG and X-gal turn the swimming surface into blue colour.

### Statistical analysis

For point-to-point comparisons in sensitivity and priming killing curves a Welch’s test was used. Growth curve analysis was carried out using the Growthcurver package for R. For survival analysis we used a log-rank test. Some specific analyses are mentioned elsewhere in the manuscript. All tests were performed with software R [81] except mutational spectrum analysis that was implemented using the *ad hoc* software iMars [48].

## Acknowledgements

We would like to thank Heidrun Häweker and Elisa Bittermann for help with library preparations and technical assistance. We are grateful to Drs Gisela Storz (NIH, Bethesda, USA) and Karen Fahrner (Harvard University, Boston, USA) for kindly supplying the strains of *E. coli* OxyS deficient mutant and the plasmid carrying FlhDC operon respectively. For mass spectrometry (M.E. and C.W.) we would like to acknowledge the assistance of the Core Facility BioSupraMol supported by the Deutsche Forschungsgemeinschaft (DFG). We also thank to Francesca Zucchetti Villagarcía and Flor I. Arias-Sánchez (from Freie Universität Berlin) for valuable help with the proofreading of the article.

## References

1. Hilker M, Schwachtje J, Baier M, Balazadeh S, Bäurle I, Geiselhardt S, et al. Priming and memory of stress responses in organisms lacking a nervous system. Biol Rev Camb Philos Soc. 2015; doi: 10.1111/brv.12215

2. Wittmann C, Chockley P, Singh SK, Pase L, Lieschke GJ, Grabher C, et al. Hydrogen Peroxide in Inflammation: Messenger, Guide, and Assassin. Adv Hematol. Hindawi Publishing Corporation; 2012;2012: 1–6. doi: 10.1155/2012/541471

3. Obata F, Fons CO, Gould AP. Early-life exposure to low-dose oxidants can increase longevity via microbiome remodelling in Drosophila. Nat Commun. Nature Publishing Group; 2018;9: 975. doi: 10.1038/s41467-018-03070-w

4. Levis NA, Pfennig DW. Evaluating ‘Plasticity-First’ Evolution in Nature: Key Criteria and Empirical Approaches. Trends Ecol Evol. 2016;31: 563–574. doi: 10.1016/j.tree.2016.03.012

5. Zheng M, Wang X, Templeton LJ, Smulski DR, LaRossa RA, Storz G. DNA microarray-mediated transcriptional profiling of the Escherichia coli response to hydrogen peroxide. J Bacteriol. 2001;183: 4562–70. doi: 10.1128/JB.183.15.4562-4570.2001

6. Zheng M, Aslund F, Storz G. Activation of the OxyR transcription factor by reversible disulfide bond formation. Science. 1998;279: 1718–21. Available: http://www.ncbi.nlm.nih.gov/pubmed/9497290

7. Imlay J a. Cellular defenses against superoxide and hydrogen peroxide. Annu Rev Biochem. 2008;77: 755–76. doi: 10.1146/annurev.biochem.77.061606.161055

8. Ezraty B, Vergnes A, Banzhaf M, Duverger Y, Huguenot A, Brochado AR, et al. Fe-S cluster biosynthesis controls uptake of aminoglycosides in a ROS-less death pathway. Science. 2013;340: 1583–7. doi: 10.1126/science.1238328

9. Imlay JA. The molecular mechanisms and physiological consequences of oxidative stress: Lessons from a model bacterium. Nature Reviews Microbiology. 2013. pp. 443–454. doi: 10.1038/nrmicro3032

10. Imlay JA. Transcription Factors That Defend Bacteria Against Reactive Oxygen Species. Annu Rev Microbiol. Annual Reviews; 2015;69: 93–108. doi: 10.1146/annurev-micro-091014-104322

11. Imlay JA, Chin SM, Linn S. Toxic DNA damage by hydrogen peroxide through the Fenton reaction in vivo and in vitro. Science (80-). 1988/04/29. 1988;240: 640–642. doi: 10.1126/science.2834821

12. Luo Y, Henle ES, Linn S. Nucleic Acids, Protein Synthesis, and Molecular Genetics: Oxidative Damage to DNA Constituents by Iron-mediated Fenton Reactions: THE DEOXYCYTIDINE FAMILY Oxidative Damage to DNA Constituents by Iron-mediated. J Biol Chem. 1996;271: 21167–76. doi: 10.1074/jbc.271.35.21167

13. Jang S, Imlay JA. Hydrogen peroxide inactivates the Escherichia coli Isc iron-sulphur assembly system, and OxyR induces the Suf system to compensate. Mol Microbiol. 2010;78: 1448–1467. doi: 10.1111/j.1365-2958.2010.07418.x

14. Imlay JA, Linn S. Bimodal pattern of killing of DNA-repair-defective or anoxically grown Escherichia coli by hydrogen peroxide. J Bacteriol. 1986;166: 519–27. Available: http://www.ncbi.nlm.nih.gov/pubmed/3516975

15. Jenkins DE, Schultz JE, Matin A. Starvation-induced cross protection against heat or H2O2 challenge in Escherichia coli. J Bacteriol. 1988;170: 3910–3914. doi: 10.1128/jb.170.9.3910-3914.1988

16. Lange R, Hengge-Aronis R. Identification of a central regulator of stationary-phase gene expression in *Escherichia coli*. Mol Microbiol. 1991;5: 49–59. doi: 10.1111/j.1365-2958.1991.tb01825.x

17. Ritz C, Baty F, Streibig JC, Gerhard D. Dose-Response Analysis Using R. Xia Y, editor. PLoS One. Public Library of Science; 2015;10: e0146021. doi: 10.1371/journal.pone.0146021

18. Basan M, Hui S, Okano H, Zhang Z, Shen Y, Williamson JR, et al. Overflow metabolism in Escherichia coli results from efficient proteome allocation. Nature. Nature Publishing Group; 2015;528: 99–104. doi: 10.1038/nature15765

19. Halliwell B, Clement MV, Long LH. Hydrogen peroxide in the human body. FEBS Lett. No longer published by Elsevier; 2000;486: 10–13. doi: 10.1016/S0014-5793(00)02197-9

20. Foyer CH, Noctor G. Stress-triggered redox signalling: What’s in pROSpect? Plant Cell and Environment. Blackwell Publishing Ltd; 2016. pp. 951–964. doi: 10.1111/pce.12621

21. Ronin I, Katsowich N, Rosenshine I, Balaban NQ. A long-term epigenetic memory switch controls bacterial virulence bimodality. Elife. 2017;6. doi: 10.7554/eLife.19599

22. Veening J-W, Smits WK, Kuipers OP. Bistability, Epigenetics, and Bet-Hedging in Bacteria. Annu Rev Microbiol. Annual Reviews; 2008;62: 193–210. doi: 10.1146/annurev.micro.62.081307.163002

23. Szklarczyk D, Franceschini A, Wyder S, Forslund K, Heller D, Huerta-Cepas J, et al. STRING v10: protein–protein interaction networks, integrated over the tree of life. Nucleic Acids Res. 2015;43: D447–D452. doi: 10.1093/nar/gku1003

24. Arifuzzaman M, Maeda M, Itoh A, Nishikata K, Takita C, Saito R, et al. Large-scale identification of protein-protein interaction of Escherichia coli K-12. Genome Res. 2006;16: 686–691. doi: 10.1101/gr.4527806

25. Basineni SR, Madhugiri R, Kolmsee T, Hengge R, Klug G. The influence of Hfq and ribonucleases on the stability of the small non-coding RNA OxyS and its target rpoS in E. coli is growth phase dependent. RNA Biol. 2009/12/18. 2009;6: 584–594. doi: 10082 [pii]

26. González-Flecha B, Demple B. Role for the oxyS gene in regulation of intracellular hydrogen peroxide in Escherichia coli. J Bacteriol. American Society for Microbiology; 1999;181: 3833–6. Available: http://www.pubmedcentral.nih.gov/articlerender.fcgi?artid=93864&tool=pmcentrez&rendertype=abstract

27. Zhang A, Altuvia S, Tiwari A, Argaman L, Hengge-Aronis R, Storz G. The OxyS regulatory RNA represses rpoS translation and binds the Hfq (HF-I) protein. EMBO J. 1998;17: 6061–6068. doi: 10.1093/emboj/17.20.6061

28. Altuvia S, Weinstein-Fischer D, Zhang A, Postow L, Storz G. A small, stable RNA induced by oxidative stress: Role as a pleiotropic regulator and antimutator. Cell. Cell Press; 1997;90: 43–53. doi: 10.1016/S0092-8674(00)80312-8

29. Barshishat S, Elgrably-Weiss M, Edelstein J, Georg J, Govindarajan S, Haviv M, et al. OxyS small RNA induces cell cycle arrest to allow DNA damage repair. EMBO J. 2018;37: 413–426. doi: 10.15252/embj.201797651

30. Willi J, Küpfer P, Eviquoz D, Fernandez G, Katz A, Leumann C, et al. Oxidative stress damages rRNA inside the ribosome and differentially affects the catalytic center. Nucleic Acids Res. Oxford University Press; 2018;46: 1945–1957. doi: 10.1093/nar/gkx1308

31. Aslund F, Zheng M, Beckwith J, Storz G. Regulation of the OxyR transcription factor by hydrogen peroxide and the cellular thiol-disulfide status. Proc Natl Acad Sci U S A. 1999;96: 6161–5. Available: http://www.ncbi.nlm.nih.gov/pubmed/10339558

32. Liu Y, Bauer SC, Imlay JA. The YaaA protein of the Escherichia coli OxyR regulon lessens hydrogen peroxide toxicity by diminishing the amount of intracellular unincorporated iron. J Bacteriol. American Society for Microbiology (ASM); 2011;193: 2186–96. doi: 10.1128/JB.00001-11

33. Ollagnier-De Choudens S, Sanakis Y, Hewitson KS, Roach P, Baldwin JE, Münck E, et al. Iron-sulfur center of biotin synthase and lipoate synthase. Biochemistry. 2000;39: 4165–73. Available: http://www.ncbi.nlm.nih.gov/pubmed/10747808

34. Hong Y, Zeng J, Wang X, Drlica K, Zhao X. Post-stress bacterial cell death mediated by reactive oxygen species. Proc Natl Acad Sci U S A. National Academy of Sciences; 2019;116: 10064–10071. doi: 10.1073/pnas.1901730116

35. Beloin C, Roux A, Ghigo JM. Escherichia coli biofilms. Current Topics in Microbiology and Immunology. 2008. pp. 249–289. doi: 10.1007/978-3-540-75418-3_12

36. Kitagawa M, Ara T, Arifuzzaman M, Ioka-Nakamichi T, Inamoto E, Toyonaga H, et al. Complete set of ORF clones of Escherichia coli ASKA library (a complete set of E. coli K-12 ORF archive): unique resources for biological research. DNA Res. 2006/06/14. 2005;12: 291–299. doi: 10.1093/dnares/dsi012

37. Pratt LA, Kolter R. Genetic analysis of *Escherichia coli* biofilm formation: roles of flagella, motility, chemotaxis and type I pili. Mol Microbiol. 1998;30: 285–293. doi: 10.1046/j.1365-2958.1998.01061.x

38. Schembri MA, Klemm P. Coordinate gene regulation by fimbriae-induced signal transduction. EMBO J. 2001;20: 3074–3081. doi: 10.1093/emboj/20.12.3074

39. Stentebjerg-Olesen B, Chakraborty T, Klemm P. FimE-catalyzed off-to-on inversion of the type 1 fimbrial phase switch and insertion sequence recruitment in an *Escherichia coli* K-12 *fimB* strain. FEMS Microbiol Lett. 2000;182: 319–325. doi: 10.1111/j.1574-6968.2000.tb08915.x

40. Avalos Vizcarra I, Hosseini V, Kollmannsberger P, Meier S, Weber SS, Arnoldini M, et al. How type 1 fimbriae help Escherichia coli to evade extracellular antibiotics. Sci Rep. Nature Publishing Group; 2016;6. doi: 10.1038/srep18109

41. Keith BR, Harris SL, Russell PW, Orndorff PE. Effect of type 1 piliation on in vitro killing of Escherichia coli by mouse peritoneal macrophages. Infect Immun. 1990;58: 3448–54. Available: http://www.ncbi.nlm.nih.gov/pubmed/1976116

42. Slauch JM. How does the oxidative burst of macrophages kill bacteria? Still an open question. Mol Microbiol. 2011;80: 580–583. doi: 10.1111/j.1365-2958.2011.07612.x

43. Wang S, Fleming RT, Westbrook EM, Matsumura P, McKay DB. Structure of the Escherichia coli FlhDC complex, a prokaryotic heteromeric regulator of transcription. J Mol Biol. 2006;355: 798–808. doi: 10.1016/j.jmb.2005.11.020

44. Barker CS, Prüss BM, Matsumura P. Increased motility of Escherichia coli by insertion sequence element integration into the regulatory region of the flhD operon. J Bacteriol. 2004;186: 7529–37. doi: 10.1128/JB.186.22.7529-7537.2004

45. Martínez-García E, Nikel PI, Chavarría M, de Lorenzo V. The metabolic cost of flagellar motion in Pseudomonas putida KT2440. Environ Microbiol. 2014;16: 291–303. doi: 10.1111/1462-2920.12309

46. Sarenko O, Klauck G, Wilke FM, Pfiffer V, Richter AM, Herbst S, et al. More than Enzymes That Make or Break Cyclic Di-GMP-Local Signaling in the Interactome of GGDEF/EAL Domain Proteins of Escherichia coli. doi: 10.1128/mBio.01639-17

47. Wang X, Kim Y, Ma Q, Hong SH, Pokusaeva K, Sturino JM, et al. Cryptic prophages help bacteria cope with adverse environments. Nat Commun. Nature Publishing Group; 2010;1: 147. doi: 10.1038/ncomms1146

48. Morgan C, Lewis PD. iMARS - Mutation analysis reporting software: An analysis of spontaneous cII mutation spectra. Mutat Res - Genet Toxicol Environ Mutagen. 2006;603: 15–26. doi: 10.1016/j.mrgentox.2005.09.010

49. Kino K, Sugiyama H. Possible cause of G·C→C·G transversion mutation by guanine oxidation product, imidazolone. Chem Biol. Cell Press; 2001;8: 369–378. doi: 10.1016/S1074-5521(01)00019-9

50. Rodríguez-Rojas A, Makarova O, Müller U, Rolff J. Cationic Peptides Facilitate Iron-induced Mutagenesis in Bacteria. Matic I, editor. Public Library of Science; 2015;11: e1005546. doi: 10.1371/journal.pgen.1005546

51. Rodríguez-Rojas A, Makarova O, Rolff J. Antimicrobials, stress and mutagenesis. Zasloff M, editor. PLoS Pathog. Public Library of Science; 2014;10: e1004445. doi: 10.1371/journal.ppat.1004445

52. Jang S, Imlay JA. Micromolar intracellular hydrogen peroxide disrupts metabolism by damaging iron-sulfur enzymes. J Biol Chem. 2006/11/15. 2007;282: 929–937. doi: M607646200 [pii] 10.1074/jbc.M607646200

53. Galhardo RS, Hastings PJ, Rosenberg SM. Mutation as a Stress Response and the Regulation of Evolvability. Crit Rev Biochem Mol Biol. 2007;42: 399–435. doi: 10.1080/10409230701648502

54. Blázquez J, Couce A, Rodríguez-Beltrán J, Rodríguez-Rojas A, Blazquez J, Couce A, et al. Antimicrobials as promoters of genetic variation. Curr Opin Microbiol. 2012/08/15. 2012;15: 561–569. doi: S1369-5274(12)00102-6 [pii]10.1016/j.mib.2012.07.007

55. Ram Y, Hadany L. THE EVOLUTION OF STRESS-INDUCED HYPERMUTATION IN ASEXUAL POPULATIONS. Evolution (N Y). 2012;66: 2315–2328. doi: 10.1111/j.1558-5646.2012.01576.x

56. Couce A, Rodríguez-Rojas A, Blázquez J. Bypass of genetic constraints during mutator evolution to antibiotic resistance. Proc Biol Sci. 2015;282: 20142698. doi: 10.1098/rspb.2014.2698

57. Datsenko KA, Wanner BL. One-step inactivation of chromosomal genes in Escherichia coli K-12 using PCR products. Proc Natl Acad Sci U S A. 2000/06/01. 2000;97: 6640–6645. doi: 10.1073/pnas.120163297 120163297 [pii]

58. Baba T, Ara T, Hasegawa M, Takai Y, Okumura Y, Baba M, et al. Construction of Escherichia coli K-12 inframe, single-gene knockout mutants: the Keio collection. Mol Syst Biol. 2006/06/02. 2006;2: 2006 0008. doi:msb4100050 [pii] 10.1038/msb4100050

59. Thomason LC, Costantino N, Court DL. *E. coli* Genome Manipulation by P1 Transduction. Current Protocols in Molecular Biology. Hoboken, NJ, USA: John Wiley & Sons, Inc.; 2007. pp. 1.17.1-1.17.8. doi: 10.1002/0471142727.mb0117s79

60. Navarro JA, Roncel M, De la Rosa FF, De la Rosa MA. Light-driven hydrogen peroxide production as a way to solar energy conversion. Bioelectrochemistry Bioenerg. Elsevier; 1987;18: 71–78. doi: 10.1016/0302-4598(87)85009-2

61. Sprouffske K, Wagner A. Growthcurver: an R package for obtaining interpretable metrics from microbial growth curves. BMC Bioinformatics. BioMed Central; 2016;17: 172. doi: 10.1186/s12859-016-1016-7

62. Rappsilber J, Mann M, Ishihama Y. Protocol for micro-purification, enrichment, pre-fractionation and storage of peptides for proteomics using StageTips. Nat Protoc. 2007;2: 1896–906. doi: 10.1038/nprot.2007.261

63. Tyanova S, Temu T, Cox J. The MaxQuant computational platform for mass spectrometry-based shotgun proteomics. Nat Protoc. Nature Publishing Group; 2016;11: 2301–2319. doi: 10.1038/nprot.2016.136

64. Tyanova S, Temu T, Sinitcyn P, Carlson A, Hein MY, Geiger T, et al. The Perseus computational platform for comprehensive analysis of (prote)omics data. Nat Methods. 2016;13: 731–40. doi: 10.1038/nmeth.3901

65. Bachmair A, Finley D, Varshavsky A. In vivo half-life of a protein is a function of its amino-terminal residue. Science (80-). 1986;234: 179–186. doi: 10.1126/science.3018930

66. Ciechanover A, Schwartz AL. How are substrates recognized by the ubiquitin-mediated proteolytic system? Trends Biochem Sci. 1989;14: 483–8. doi: 10.1016/0968-0004(89)90180-1

67. Tobias JW, Shrader TE, Rocap G, Varshavsky A. The N-end rule in bacteria. Science. 1991;254: 1374–7. doi: 10.1126/science.1962196

68. Guruprasad K, Reddy B V, Pandit MW. Correlation between stability of a protein and its dipeptide composition: a novel approach for predicting in vivo stability of a protein from its primary sequence. Protein Eng. 1990;4: 155–61. doi: 10.1093/protein/4.2.155

69. Wilkins MR, Gasteiger E, Bairoch A, Sanchez J-C, Williams KL, Appel RD, et al. Protein Identification and Analysis Tools in the ExPASy Server. 2-D Proteome Analysis Protocols. New Jersey: Humana Press; 1999. pp. 531–552. doi: 10.1385/1-59259-584-7:531

70. Humbard MA, Surkov S, De Donatis GM, Jenkins LM, Maurizi MR. The N-degradome of Escherichia coli: limited proteolysis in vivo generates a large pool of proteins bearing N-degrons. J Biol Chem. American Society for Biochemistry and Molecular Biology; 2013;288: 28913–24. doi: 10.1074/jbc.M113.492108

71. Pfaffl MW, Tichopad A, Prgomet C, Neuvians TP. Determination of stable housekeeping genes, differentially regulated target genes and sample integrity: BestKeeper--Excel-based tool using pair-wise correlations. Biotechnol Lett. 2004;26: 509–15. Available: http://www.ncbi.nlm.nih.gov/pubmed/15127793

72. Pfaffl MW, Horgan GW, Dempfle L. Relative expression software tool (REST) for group-wise comparison and statistical analysis of relative expression results in real-time PCR. Nucleic Acids Res. 2002;30: e36. Available: http://www.pubmedcentral.nih.gov/articlerender.fcgi?artid=113859&tool=pmcentrez&rendertype=abstract

73. Bustin SA, Benes V, Garson JA, Hellemans J, Huggett J, Kubista M, et al. The MIQE guidelines: minimum information for publication of quantitative real-time PCR experiments. Clin Chem. 2009;55: 611–22. doi: 10.1373/clinchem.2008.112797

74. Seemann T. Snippy: fast bacterial variant calling from NGS reads [Internet]. 2015. Available: https://github.com/tseemann/snippy

75. Li H, Durbin R. Fast and accurate short read alignment with Burrows-Wheeler transform. Bioinformatics. 2009;25: 1754–1760. doi: 10.1093/bioinformatics/btp324

76. Garrison E, Marth G. Haplotype-based variant detection from short-read sequencing. 2012; Available: https://arxiv.org/pdf/1207.3907.pdf

77. Deatherage DE, Barrick JE. Identification of Mutations in Laboratory-Evolved Microbes from Next-Generation Sequencing Data Using breseq. Methods in molecular biology (Clifton, NJ). 2014. pp. 165–188. doi: 10.1007/978-1-4939-0554-6_12

78. Sanders LH, Sudhakaran J, Sutton MD. The GO system prevents ROS-induced mutagenesis and killing in Pseudomonas aeruginosa. FEMS Microbiol Lett. 2009/03/18. 2009;294: 89–96. doi: FML1550 [pii] 10.1111/j.1574-6968.2009.01550.x

79. Rodriguez-Rojas A, Blazquez J, Rodríguez-Rojas A, Blázquez J. The Pseudomonas aeruginosa pfpI gene plays an antimutator role and provides general stress protection. J Bacteriol. 2008/11/26. American Society for Microbiology; 2009;191: 844–850. doi: JB.01081-08 [pii] 10.1128/JB.01081-08

80. Fahrner KA, Berg HC. Mutations That Stimulate flhDC Expression in Escherichia coli K-12. J Bacteriol. 2015;197: 3087–96. doi: 10.1128/JB.00455-15

81. R Development Core Team. RStudio Team (2015). RStudio: Integrated Development for R. [Internet]. Boston: MA URL http://www.rstudio.com/; 2015. Available: http://www.rstudio.com

