## Supplemental figures and supplemental table legends for "Non-lethal exposure to H_2_O_2_ boosts bacterial survival and evolvability against oxidative stress"

A

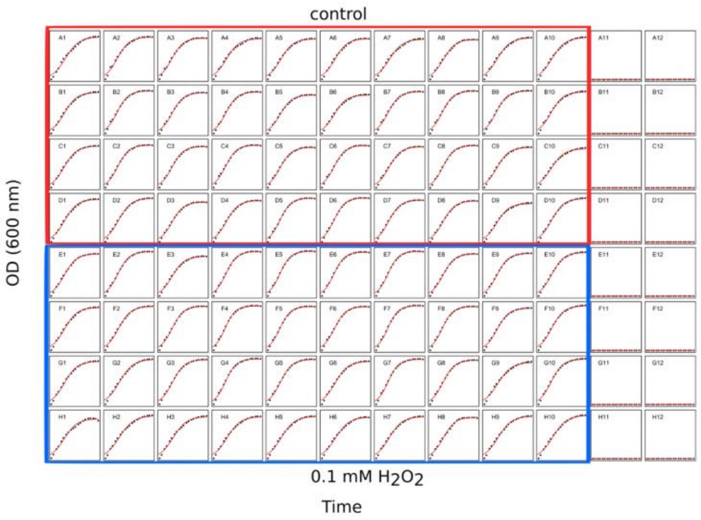


B

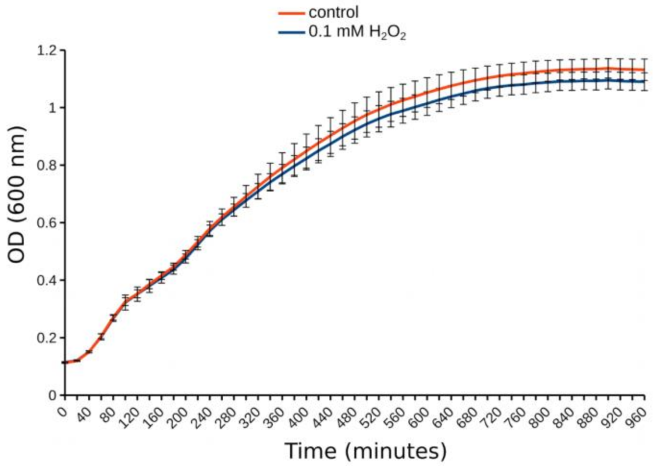


**S1 Fig. Growth curves of *E. coli* cells treated with 0.1 mM $H_2O_2$ and non-treated ones (control). Panel A represents a graphical model fitting of individual curves plotted by Growthcurver R package [60]. Panel B shows the average growth curves from 40 independent replicas per each situation (0.1 mM $H_2O_2$ versus control).**

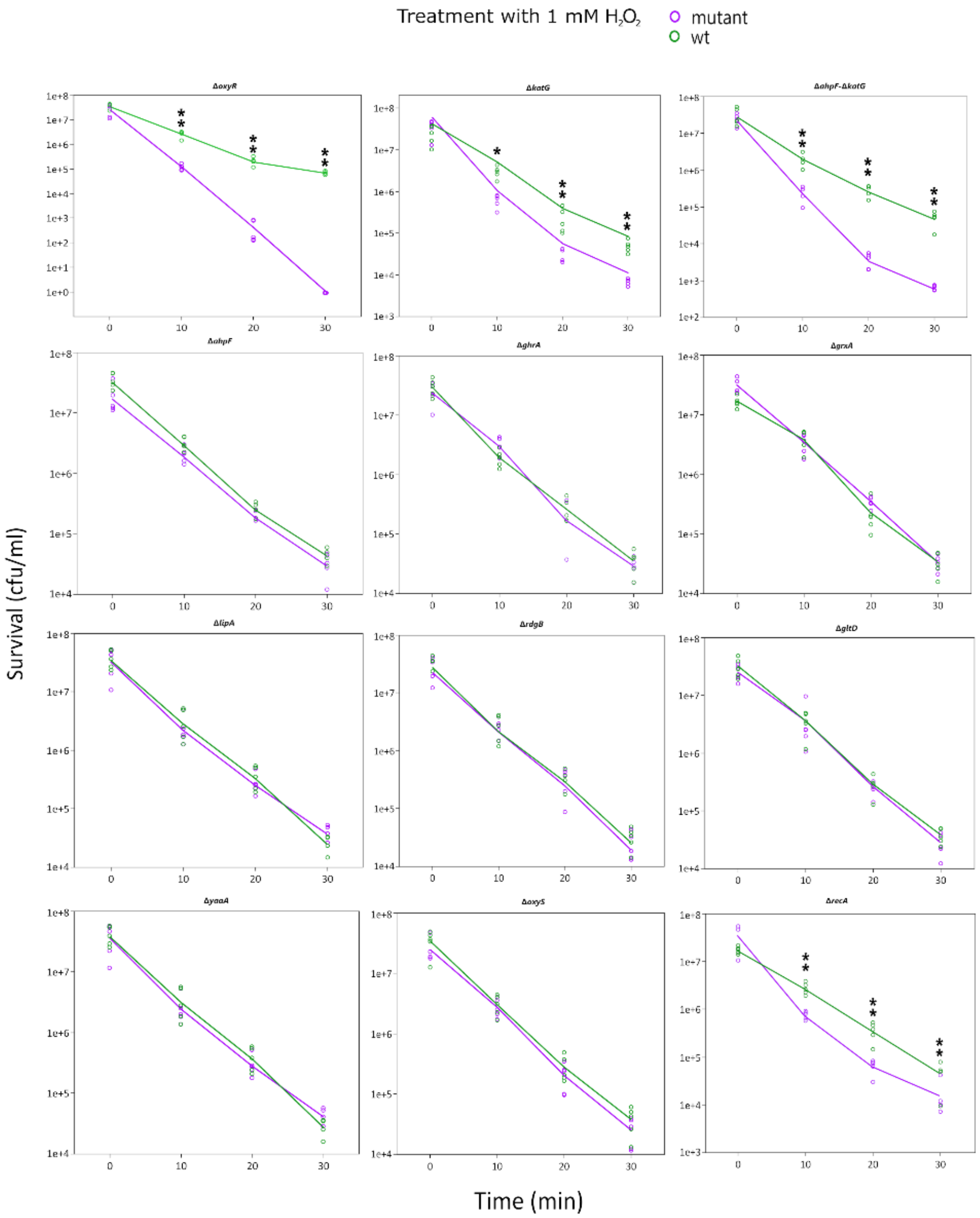


**S2 Fig. Sensitivity of a selected set of mutants in relevant genes of *E. coli* MG1655 that presented an increased level of expression as part of the priming response. Asterisks represent significant differences between the wild-type (wt) strain and its derivatives mutants (Welch's test, one asterisk for *p*<0.05 and two asterisks for *p*<0.01).**

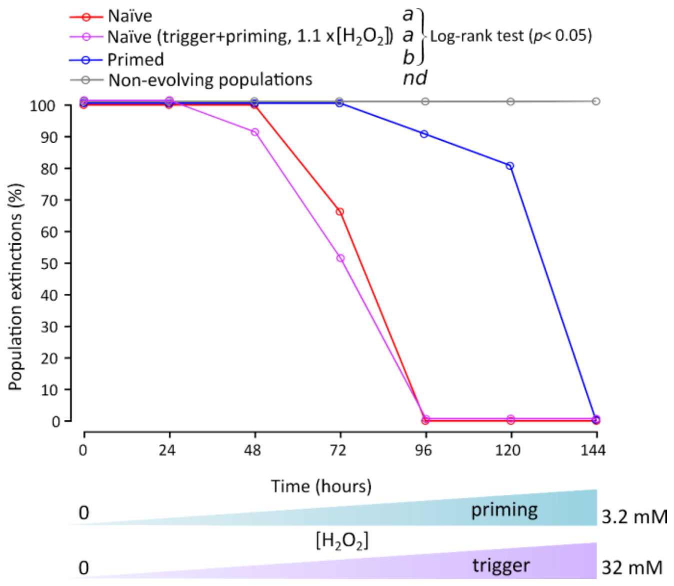


**S3 Fig. Repetition of the evolution experiment including a control treated with priming and trigger concentration simultaneously (magenta line). The extinction was perceived by negative growth in the next passage and by the absence of growth in LB plates during contamination controls. Non-evolving population control (grey line, 20 populations) is presented. Evolvability differs between the two naïve population groups (red and magenta lines) and primed populations (blue line). Equal letter represents no statistical differences while the same letter indicates significant differences in pair-wise comparison (Log-rank test, $p < 0.05$). Differences with non-evolved populations were not determined.**

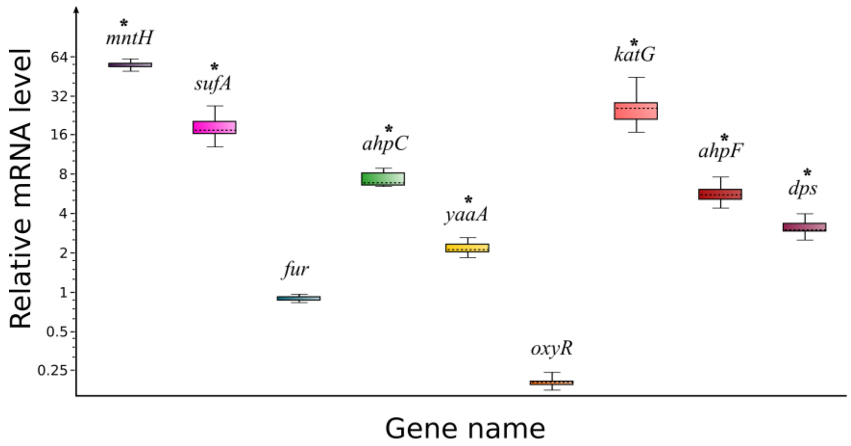


**S4 Fig. Relative gene expression for *E. coli* MG1655 OxyR regulon selected genes in treatments with 0.1 mM $H_2O_2$ versus non-treated bacteria. Cells were collected 30 minutes after exposure. Error bars represent the standard error of the mean of three independent biological replicates, each biological replicate is the average of three technical repetitions.**

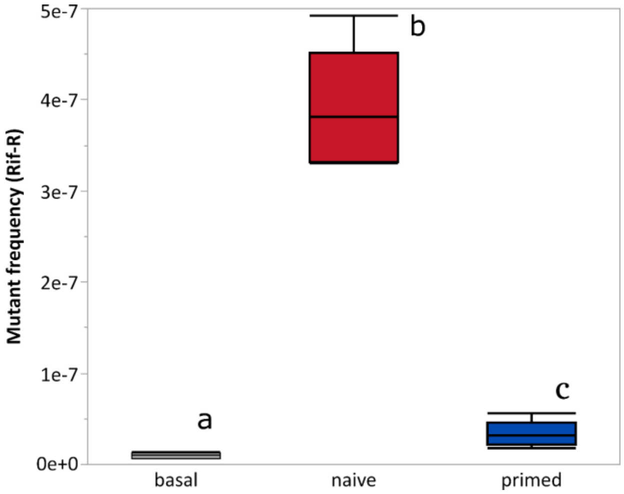


**S5 Fig. Boxplot showing the $H_2O_2$-induced mutant frequency. Naïve and primed cells (pre-treated with 0.1 mM, 30 minutes in advanced) cultures challenged with 1 mM, allowed to recover and plated in rifampicin (100 µg/ml). The basal level of mutagenesis for non-pre-treated, non-challenged cells is also shown. Every sample consisted of five independent replications. Letters denote significant differences (Welch's test, *p*=0.03 for basal level versus primed, *p*<0.01 for both basal versus naïve and primed versus naïve).**

### Supplementary table legends.

**S1 Table. Growth curve parameters showing priming costs for *E. coli* cells treated with 0.1 mM $H_2O_2$ when compared to untreated controls. The growth curve parameters were estimated with the Growthcurver R package [61]. Only carrying capacity and the areas under the curve have shown significant differences with a small effect.**

**S2 Table. Quantitative $H_2O_2$ determination of *E. coli* culture supernatants after a treatment with 1 mM after priming bacteria with 0.1 mM $H_2O_2$ in comparison to naïve cells. $H_2O_2$ concentrations were determined for 0, 15 and 30 minutes after the addition of $H_2O_2$ using the Pierce Quantitative Peroxide kit (Thermo Scientific, Germany). The shown values represent the mean of the supernatant from three individual cultures and their standard deviations.**

**S3 Table. Output table of the proteomic experiment reporting time-lapse decline of the *E. coli* response to 1 mM $H_2O_2$ for 30 minutes. Bacteria were sampled 30, 60, 90, 120 and 150 minutes after removal of the treatment. Each treatment group consisted of six independent replicates and bacteria before treatment (T0) were used as control. Statistical analysis used student t-test and false discovery rate for correction of the p-values (data analysis using Maxquant and Perseus software for label-free quantification of proteins with LC-MS).**

**S4 Table. Output table of the proteomic experiment reporting *E. coli* response to $H_2O_2$ treatment of 0.1 and 1 mM during 5 minutes. Each treatment group consisted of six independent replications and bacteria before treatment (T0) were used as control. Statistical analysis used student t-test and false discovery rate for correction of the p-values (data analysis using Maxquant and Perseus software for label-free quantification of proteins with LC-MS).**

**S5 Table. Protein stability predicted from the sequences of selected proteins that showed an elevated level of expression after treatment with 0.1 mM $H_2O_2$ (priming concentration) and remained up or declined during memory duration of priming response. The predictions were carried out using the online tool ProtParam [69].**

**S6 Table. Transcripts differentially expressed (±2 log2) in the fraction of small of RNA (<200 nt) during the decay of $H_2O_2$ response (120 minutes after removal the treatment).**

**S7 Table. Transcripts differentially expressed (± 2.5 log2) in the fraction of large RNA (>200 nt) during the decay of $H_2O_2$ response (120 minutes after removal the treatment).**

**S8 Table. Relative gene expression (qPCR) for *E. coli* MG1655 selected responsive genes after a treatment with 0.1 mM $H_2O_2$ versus non-treated bacteria during 30 minutes.**

**S9 Table. Strains used in this work and their relevant phenotypes.**

**S10 Table. Primers used for amplification of kanamycin insertion of mutants from the Keio collection which PCR product was used to transfer the mutations to the *E. coli* MG1655 strain.**

**S11 Table. Primers used for relative gene expression quantification by real time PCR (qPCR).**
