## supplemental tables for "Non-lethal exposure to H_2_O_2_ boosts bacterial survival and evolvability against oxidative stress": Table S2.pdf

Table S2. Quantitative $H_2O_2$ determination of *E. coli* culture supernatants after a treatment with 1 mM after priming bacteria with 0.1 mM $H_2O_2$ in comparison to naïve cells. $H_2O_2$ concentrations were determined for 0, 15 and 30 minutes after the addition of $H_2O_2$ using the Pierce Quantitative Peroxide kit (Thermo Scientific, Germany). The shown values represent the mean of the supernatant from three individual cultures and their standard deviations.

| | Treatment [$H_2O_2$] µM (*mean*± *sd*) | | Statistical inferences | |
|---|---|---|---|---|
| Time (min) | Naive | Primed | Fold-change ratio (naive/primed) | *p*-value |
| 0 | 1062.03±44.90 | 1053.97± 69.58 | 0.99 | 0.4380 |
| 15 | 723.68±62.43 | 140.46±33.45 | 5.15 | 0.00014 |
| 30 | 379.02±49.56 | 22.37 ±23.06 | 16.94 | 0.00035 |
