## supplemental tables for "Non-lethal exposure to H_2_O_2_ boosts bacterial survival and evolvability against oxidative stress": Table S5.pdf

Table S5. Stability predicted from the sequences of selected proteins that showed an elevated level of expression after treatment with $H_2O_2$ 0.1 mM (priming concentration) and remained up or declined during memory duration of priming response.

| Protein | Estimated half-life | Stability index | Classification |
|---|---|---|---|
| KatG | >10 hours | 27.00 | stable |
| AhpC | >10 hours | 33.31 | stable |
| AhpF | >10 hours | 28.11 | stable |
| SufA | >10 hours | 40.85 | unstable |
| OxyR | >10 hours | 42.21 | unstable |
| GrxA | >10 hours | 25.29 | stable |
| GhrA | >10 hours | 38.28 | stable |
| YaaA | >10 hours | 32.05 | stable |
| Dps | >2 min | 24.99 | unstable |

Amino acid sequence evidence

KatG

MSTSDDIHNT TATGKCPFHQ GGHDQSAGAG TTTRDWWPNQ LRVDLLNQHS NRSNPLGEDF
DYRKEFSKLD YYGLKKDLKA LLTESQPWWP ADWGSYAGLF IRMAWHGAGT YRSIDGRGGA
GRGQQRFAPL NSWPDNVSLD KARRLLWPIK QKYGQKISWA DLFILAGNVA LENSGFRTFG
FGAGREDVWE PDLDVNWGDE KAWLTHRHPE ALAKAPLGAT EMGLIYVNPE GPDHSGEPLS
AAAAIRATFG NMGMNDEETV ALIAGGHTLG KTHGAGPTSN VGPDPEAAPI EEQGLGWAST
YGSGVGADAI TSGLEVVWTQ TPTQWSNYFF ENLFKYEWVQ TRSPAGAIQF EAVDAPEIIP
DPFDPSKKRK PTMLVTDLTL RFDPEFEKIS RRFLNDPQAF NEAFARAWFK LTHRDMGPKS
RYIGPEVPKE DLIWQDPLPQ PIYNPTEQDI IDLKFAIADS GLSVSELVSV AWASASTFRG
GDKRGGANGA RLALMPQRDW DVNAAAVRAL PVLEKIQKES GKASLADIIV LAGVVGVEKA

ASAAGLSIHV PFAPGRVDAR QDQTDIEMFE LLEPIADGFR NYRARLDVST TESLLIDKAQ QLTLTAPEMT ALVGGMRVLG ANFDGSKNGV FTDRVGVLSN DFFVNLLDMR YEWKATDESK
ELFEGRDRET GEVKFTASRA DLVFGSNSVL RAVAEVYASS DAHEKFVKDF VAAWVKVMNL
DRFDLL

Number of amino acids: 726

Molecular weight: 80023.82

Theoretical pI: 5.14

Amino acid composition:

Ala (A) 77 10.6%

Arg (R) 41 5.6%

Asn (N) 28 3.9%

Asp (D) 55 7.6%

Cys (C) 1 0.1%

Gln (Q) 24 3.3%

Glu (E) 44 6.1%

Gly (G) 65 9.0%

His (H) 13 1.8%

Ile (I) 28 3.9%

Leu (L) 63 8.7%

| | | |
|---|---|---|
| Lys (K) | 35 | 4.8% |
| Met (M) | 13 | 1.8% |
| Phe (F) | 36 | 5.0% |
| Pro (P) | 40 | 5.5% |
| Ser (S) | 44 | 6.1% |
| Thr (T) | 39 | 5.4% |
| Trp (W) | 22 | 3.0% |
| Tyr (Y) | 15 | 2.1% |
| Val (V) | 43 | 5.9% |
| Pyl (O) | 0 | 0.0% |
| Sec (U) | 0 | 0.0% |
| (B) | 0 | 0.0% |
| (Z) | 0 | 0.0% |
| (X) | 0 | 0.0% |

Total number of negatively charged residues (Asp + Glu): 99

Total number of positively charged residues (Arg + Lys): 76

Estimated half-life:

The N-terminal of the sequence considered is M (Met).

The estimated half-life is: 30 hours (mammalian reticulocytes, in vitro).

>20 hours (yeast, in vivo).

>10 hours (Escherichia coli, in vivo).

Instability index:

The instability index (II) is computed to be 27.00

This classifies the protein as stable.

Aliphatic index: 76.67

Grand average of hydropathicity (GRAVY): -0.372

AhpC

MSLINTKIKP FKNQAFKNGE FIEITEKDTE GRWSVFFFYP ADFTFVCPTE LGDVADHYEE LQKLGVDVYA VSTDTHFTHK AWHSSSETIA KIKYAMIGDP TGALTRNFDN MREDEGLADR ATFVVDPQGI IQAIEVTAEG IGRDASDLLR KIKAAQYVAS HPGEVCPAKW KEGEATLAPS LDLVGKI

Number of amino acids: 187

Molecular weight: 20761.44

Theoretical pI: 5.03

Amino acid composition:

| | | |
|---|---|---|
| Ala (A) | 19 | 10.2% |
| Arg (R) | 6 | 3.2% |
| Asn (N) | 5 | 2.7% |
| Asp (D) | 14 | 7.5% |
| Cys (C) | 2 | 1.1% |
| Gln (Q) | 5 | 2.7% |
| Glu (E) | 15 | 8.0% |
| Gly (G) | 13 | 7.0% |
| His (H) | 5 | 2.7% |
| Ile (I) | 13 | 7.0% |
| Leu (L) | 11 | 5.9% |
| Lys (K) | 14 | 7.5% |
| Met (M) | 3 | 1.6% |

| | | |
|---|---|---|
| Phe (F) | 11 | 5.9% |
| Pro (P) | 8 | 4.3% |
| Ser (S) | 9 | 4.8% |
| Thr (T) | 14 | 7.5% |
| Trp (W) | 3 | 1.6% |
| Tyr (Y) | 5 | 2.7% |
| Val (V) | 12 | 6.4% |
| Pyl (O) | 0 | 0.0% |
| Sec (U) | 0 | 0.0% |

Estimated half-life:

The N-terminal of the sequence considered is M (Met).

The estimated half-life is: 30 hours (mammalian reticulocytes, in vitro).

>20 hours (yeast, in vivo).

>10 hours (Escherichia coli, in vivo).

Instability index:

The instability index (II) is computed to be 33.31

This classifies the protein as stable.

Aliphatic index: 78.82

Grand average of hydropathicity (GRAVY): -0.278

AhpF

MLDTNMKTQL KAYLEKLTKP VELIATLDDS AKSAEIKELL AEIAELSDKV TFKEDNSLPV
RKPSFLITNP GSNQGPRFAG SPLGHEFTSL VLALLWTGGH PSKEAQSLLE QIRHIDGDFE
FETYYSLSCH NCPDVVQALN LMSVLNPRIK HTAIDGGTFQ NEITDRNVMG VPAVFVNGKE
FGQGRMTLTE IVAKIDTGAE KRAAEELNKR DAYDVLIVGS GPAGAAAAIY SARKGIRTGL
MGERFGGQIL DTVDIENYIS VPKTEGQKLA GALKVHVDEY DVDVIDSQSA SKLIPAAVEG
GLHQIETASG AVLKARSIIV ATGAKWRNMN VPGEDQYRTK GVTYCPHCDG PLFKGKRVAV
IGGGNSGVEA AIDLAGIVEH VTLLEFAPEM KADQVLQDKL RSLKNVDIIL NAQTTEVKGD
GSKVVGLEYR DRVSGDIHNI ELAGIFVQIG LLPNTNWLEG AVERNRMGEI IIDAKCETNV
KGVFAAGDCT TVPYKQIIIA TGEGAKASLS AFDYLIRTKT A

Number of amino acids: 521

Molecular weight: 56177.11

Theoretical pI: 5.47

Amino acid composition:

Ala (A) 50 9.6%

| | | |
|---|---|---|
| Arg (R) | 21 | 4.0% |
| Asn (N) | 22 | 4.2% |
| Asp (D) | 31 | 6.0% |
| Cys (C) | 6 | 1.2% |
| Gln (Q) | 17 | 3.3% |
| Glu (E) | 37 | 7.1% |
| Gly (G) | 49 | 9.4% |
| His (H) | 10 | 1.9% |
| Ile (I) | 38 | 7.3% |
| Leu (L) | 47 | 9.0% |
| Lys (K) | 35 | 6.7% |
| Met (M) | 9 | 1.7% |
| Phe (F) | 15 | 2.9% |
| Pro (P) | 19 | 3.6% |
| Ser (S) | 27 | 5.2% |
| Thr (T) | 32 | 6.1% |
| Trp (W) | 3 | 0.6% |
| Tyr (Y) | 12 | 2.3% |
| Val (V) | 41 | 7.9% |

Pyl (O) 0 0.0%

Sec (U) 0 0.0%

(B) 0 0.0%

(Z) 0 0.0%

(X) 0 0.0%

Total number of negatively charged residues (Asp + Glu): 68

Total number of positively charged residues (Arg + Lys): 56

Ext. coefficient 34380

Abs 0.1% (=1 g/l) 0.612, assuming all Cys residues are reduced

Estimated half-life:

The N-terminal of the sequence considered is M (Met).

The estimated half-life is: 30 hours (mammalian reticulocytes, in vitro).

>20 hours (yeast, in vivo).

>10 hours (Escherichia coli, in vivo).

Instability index:

The instability index (II) is computed to be 28.11

This classifies the protein as stable.

Aliphatic index: 96.05

Grand average of hydropathicity (GRAVY): -0.123

SufA

MDMHSGTFNP QDFAWQGLTL TPAAAIHIRE LVAKQPGMVG VRLGVKQTGC AGFGYVLDSV
SEPDKDDLLF EHDGAKLFVP LQAMPFIDGT EVDFVREGLN QIFKFHNPKA QNECGCGESF GV

Number of amino acids: 122

Molecular weight: 13300.11

Theoretical pI: 4.85

Amino acid composition:

Ala (A) 9 7.4%

Arg (R) 3 2.5%

Asn (N) 4 3.3%

| | | |
|---|---|---|
| Asp (D) | 9 | 7.4% |
| Cys (C) | 3 | 2.5% |
| Gln (Q) | 7 | 5.7% |
| Glu (E) | 7 | 5.7% |
| Gly (G) | 14 | 11.5% |
| His (H) | 4 | 3.3% |
| Ile (I) | 4 | 3.3% |
| Leu (L) | 10 | 8.2% |
| Lys (K) | 6 | 4.9% |
| Met (M) | 4 | 3.3% |
| Phe (F) | 10 | 8.2% |
| Pro (P) | 7 | 5.7% |
| Ser (S) | 4 | 3.3% |
| Thr (T) | 5 | 4.1% |
| Trp (W) | 1 | 0.8% |
| Tyr (Y) | 1 | 0.8% |
| Val (V) | 10 | 8.2% |
| Pyl (O) | 0 | 0.0% |
| Sec (U) | 0 | 0.0% |

| | | |
|---|---|---|
| (B) | 0 | 0.0% |
| (Z) | 0 | 0.0% |
| (X) | 0 | 0.0% |

Total number of negatively charged residues (Asp + Glu): 16

Total number of positively charged residues (Arg + Lys): 9

Ext. coefficient 6990

Abs 0.1% (=1 g/l) 0.526, assuming all Cys residues are reduced

Estimated half-life:

The N-terminal of the sequence considered is M (Met).

The estimated half-life is: 30 hours (mammalian reticulocytes, in vitro).

>20 hours (yeast, in vivo).

>10 hours (Escherichia coli, in vivo).

Instability index:

The instability index (II) is computed to be 40.85

This classifies the protein as unstable.

Aliphatic index: 75.90

Grand average of hydropathicity (GRAVY): -0.103

OxyR

MNIRDLEYLV ALAEHRHFRR AADSCHVSQP TLSGQIRKLE DELGVMLLER TSRKVLFTQA
GMLLVDQART VLREVKVLKE MASQQGETMS GPLHIGLIPT VGPYLLPHII PMLHQTFPKL
EMYLHEAQTH QLLAQLDSGK LDCVILALVK ESEAFIEVPL FDEPMLLAIY EDHPWANREC
VPMADLAGEK LLMLEDGHCL RDQAMGFCFE AGADEDTHFR ATSLETLRNM VAAGSGITLL
PALAVPPERK RDGVVYLPCI KPEPRRTIGL VYRPGSPLRS RYEQLAEAIR ARMDGHFDKV
LKQAV

Number of amino acids: 305

Molecular weight: 34275.93

Theoretical pI: 5.96

Amino acid composition:

| | | |
|---|---|---|
| Ala (A) | 27 | 8.9% |
| Arg (R) | 22 | 7.2% |
| Asn (N) | 3 | 1.0% |
| Asp (D) | 16 | 5.2% |
| Cys (C) | 6 | 2.0% |
| Gln (Q) | 13 | 4.3% |
| Glu (E) | 25 | 8.2% |
| Gly (G) | 18 | 5.9% |
| His (H) | 12 | 3.9% |
| Ile (I) | 13 | 4.3% |
| Leu (L) | 44 | 14.4% |
| Lys (K) | 12 | 3.9% |
| Met (M) | 13 | 4.3% |
| Phe (F) | 9 | 3.0% |
| Pro (P) | 19 | 6.2% |

Ser (S) 12 3.9%

Thr (T) 13 4.3%

Trp (W) 1 0.3%

Tyr (Y) 7 2.3%

Val (V) 20 6.6%

Pyl (O) 0 0.0%

Sec (U) 0 0.0%

(B) 0 0.0%

(Z) 0 0.0%

(X) 0 0.0%

Total number of negatively charged residues (Asp + Glu): 41

Total number of positively charged residues (Arg + Lys): 34

Estimated half-life:

The N-terminal of the sequence considered is M (Met).

The estimated half-life is: 30 hours (mammalian reticulocytes, in vitro).

>20 hours (yeast, in vivo).

>10 hours (Escherichia coli, in vivo).

Instability index:

The instability index (II) is computed to be 42.21

This classifies the protein as unstable.

Aliphatic index: 100.75

Grand average of hydropathicity (GRAVY): -0.088

GrxA

MQTVIFGRSG CPYCVRAKDL AEKLSNERDD FQYQYVDIRA EGITKEDLQQ KAGKPVETVP
QIFVDQQHIG GYTDFAAWVK ENLDA

**Number of amino acids:** 85

**Molecular weight:** 9684.85

**Theoretical pI:** 4.81

Amino acid composition:

| | | |
|---|---|---|
| Ala (A) | 7 | 8.2% |
| Arg (R) | 4 | 4.7% |
| Asn (N) | 2 | 2.4% |
| Asp (D) | 8 | 9.4% |
| Cys (C) | 2 | 2.4% |
| Gln (Q) | 8 | 9.4% |
| Glu (E) | 6 | 7.1% |
| Gly (G) | 6 | 7.1% |
| His (H) | 1 | 1.2% |
| Ile (I) | 5 | 5.9% |
| Leu (L) | 4 | 4.7% |
| Lys (K) | 6 | 7.1% |
| Met (M) | 1 | 1.2% |
| Phe (F) | 4 | 4.7% |
| Pro (P) | 3 | 3.5% |
| Ser (S) | 2 | 2.4% |

| | | |
|---|---|---|
| Thr (T) | 4 | 4.7% |
| Trp (W) | 1 | 1.2% |
| Tyr (Y) | 4 | 4.7% |
| Val (V) | 7 | 8.2% |
| Pyl (O) | 0 | 0.0% |
| Sec (U) | 0 | 0.0% |
| (B) | 0 | 0.0% |
| (Z) | 0 | 0.0% |
| (X) | 0 | 0.0% |

Total number of negatively charged residues (Asp + Glu): 14

Total number of positively charged residues (Arg + Lys): 10

Estimated half-life:

The N-terminal of the sequence considered is M (Met).

The estimated half-life is: 30 hours (mammalian reticulocytes, in vitro).

>20 hours (yeast, in vivo).

>10 hours (Escherichia coli, in vivo).

Instability index:

The instability index (II) is computed to be 25.29

This classifies the protein as stable.

Aliphatic index: 73.41

Grand average of hydropathicity (GRAVY): -0.571

GhrA

MDIIFYHPTF DTQWWIEALR KAIPQARVRA WKSGDNDSAD YALVWHPPVE MLAGRDLKAV
FALGAGVDSI LSKLQAHPEM LNPSVPLFRL EDTGMGEQMQ EYAVSQVLHW FRRFDDYRIQ
QNSSHWQPLP EYHREDFTIG ILGAGVLGSK VAQSLQTWRF PLRCWSRTRK SWPGVQSFAG
REELSAFLSQ CRVLINLLPN TPETVGIINQ QLLEKLPDGA YLLNLARGVH VVEDDLLAAL
DSGKVKGAML DVFNREPLPP ESPLWQHPRV TITPHVAAIT RPAEAVEYIS RTIAQLEKGE
RVCGQVDRAR GY

Number of amino acids: 312

Molecular weight: 35343.42

Theoretical pI: 6.32

Amino acid composition:

Ala (A) 27 8.7%

Arg (R) 23 7.4%

Asn (N) 8 2.6%

Asp (D) 17 5.4%

Cys (C) 3 1.0%

Gln (Q) 18 5.8%

Glu (E) 19 6.1%

Gly (G) 20 6.4%

His (H) 9 2.9%

Ile (I) 15 4.8%

Leu (L) 34 10.9%

Lys (K) 10 3.2%

Met (M) 6 1.9%

Phe (F) 11 3.5%

Pro (P) 21 6.7%

| | | |
|---|---|---|
| Ser (S) | 18 | 5.8% |
| Thr (T) | 12 | 3.8% |
| Trp (W) | 10 | 3.2% |
| Tyr (Y) | 8 | 2.6% |
| Val (V) | 23 | 7.4% |
| Pyl (O) | 0 | 0.0% |
| Sec (U) | 0 | 0.0% |

| | | |
|---|---|---|
| (B) | 0 | 0.0% |
| (Z) | 0 | 0.0% |
| (X) | 0 | 0.0% |

Total number of negatively charged residues (Asp + Glu): 36

Total number of positively charged residues (Arg + Lys): 33

Estimated half-life:

The N-terminal of the sequence considered is M (Met).

The estimated half-life is: 30 hours (mammalian reticulocytes, in vitro).

>20 hours (yeast, in vivo).

>10 hours (Escherichia coli, in vivo).

Instability index:

The instability index (II) is computed to be 38.28

This classifies the protein as stable.

Aliphatic index: 91.28

Grand average of hydropathicity (GRAVY): -0.258

YaaA

MLILISPAKT LDYQSPLTTT RYTLPELLDN SQQLIHEARK LTPPQISTLM RISDKLAGIN
AARFHDWQPD FTPANARQAI LAFKGDVYTG LQAETFSEDD FDFAQQHLRM LSGLYGVLRP
LDLMQPYRLE MGIRLENARG KDLYQFWGDI ITNKLNEALA AQGDNVVINL ASDEYFKSVK
PKKLNAEIIK PVFLDEKNGK FKIISFYAKK ARGLMSRFII ENRLTKPEQL TGFNSEGYFF
DEDSSSNGEL VFKRYEQR

Number of amino acids: 258

Molecular weight: 29585.83

Theoretical pI: 6.86

Amino acid composition:

| | | |
|---|---|---|
| Ala (A) | 19 | 7.4% |
| Arg (R) | 15 | 5.8% |
| Asn (N) | 13 | 5.0% |
| Asp (D) | 17 | 6.6% |
| Cys (C) | 0 | 0.0% |
| Gln (Q) | 14 | 5.4% |
| Glu (E) | 16 | 6.2% |
| Gly (G) | 14 | 5.4% |
| His (H) | 3 | 1.2% |
| Ile (I) | 17 | 6.6% |
| Leu (L) | 31 | 12.0% |
| Lys (K) | 18 | 7.0% |
| Met (M) | 6 | 2.3% |
| Phe (F) | 16 | 6.2% |
| Pro (P) | 12 | 4.7% |
| Ser (S) | 15 | 5.8% |

| | | |
|---|---|---|
| Thr (T) | 13 | 5.0% |
| Trp (W) | 2 | 0.8% |
| Tyr (Y) | 10 | 3.9% |
| Val (V) | 7 | 2.7% |
| Pyl (O) | 0 | 0.0% |
| Sec (U) | 0 | 0.0% |
| (B) | 0 | 0.0% |
| (Z) | 0 | 0.0% |
| (X) | 0 | 0.0% |

Total number of negatively charged residues (Asp + Glu): 33

Total number of positively charged residues (Arg + Lys): 33

Estimated half-life:

The N-terminal of the sequence considered is M (Met).

The estimated half-life is: 30 hours (mammalian reticulocytes, in vitro).

>20 hours (yeast, in vivo).

>10 hours (Escherichia coli, in vivo).

Instability index:

The instability index (II) is computed to be 32.05

This classifies the protein as stable.

Aliphatic index: 87.79

Grand average of hydropathicity (GRAVY): -0.403

Dps

MSTAKLVKSK ATNLLYTRND VSDSEKKATV ELLNRQVIQF IDLSLITKQA HWNMRGANFI
AVHEMLDGFR TALIDHLDTM AERAVQLGGV ALGTTQVINS KTPLKSYPLD IHNVQDHLKE
LADRYAIVAN DVRKAIGEAK DDDTADILTA ASRDLDKFLW FIESNIE

Number of amino acids: 166

Molecular weight: 18564.11

Theoretical pI: 5.70

Amino acid composition:

| | | |
|---|---|---|
| Ala (A) | 18 | 10.8% |
| Arg (R) | 8 | 4.8% |
| Asn (N) | 9 | 5.4% |
| Asp (D) | 16 | 9.6% |
| Cys (C) | 0 | 0.0% |
| Gln (Q) | 6 | 3.6% |

| | | |
|---|---|---|
| Glu (E) | 8 | 4.8% |
| Gly (G) | 6 | 3.6% |
| His (H) | 5 | 3.0% |
| Ile (I) | 12 | 7.2% |
| Leu (L) | 19 | 11.4% |
| Lys (K) | 12 | 7.2% |
| Met (M) | 3 | 1.8% |
| Phe (F) | 5 | 3.0% |
| Pro (P) | 2 | 1.2% |
| Ser (S) | 9 | 5.4% |
| Thr (T) | 12 | 7.2% |
| Trp (W) | 2 | 1.2% |
| Tyr (Y) | 3 | 1.8% |
| Val (V) | 11 | 6.6% |
| Pyl (O) | 0 | 0.0% |
| Sec (U) | 0 | 0.0% |

| | | |
|---|---|---|
| (B) | 0 | 0.0% |
| (Z) | 0 | 0.0% |
| (X) | 0 | 0.0% |

Total number of negatively charged residues (Asp + Glu): 24
Total number of positively charged residues (Arg + Lys): 20

Atomic composition:

| | | |
|---|---|---|
| Carbon | C | 821 |
| Hydrogen | H | 1325 |
| Nitrogen | N | 229 |
| Oxygen | O | 254 |
| Sulfur | S | 3 |

Formula: C821H1325N229O254S3
Total number of atoms: 2632

Extinction coefficients:

Extinction coefficients are in units of M-1 cm-1, at 280 nm measured in water.

Ext. coefficient 15470
Abs 0.1% (=1 g/l) 0.833

Estimated half-life:

The N-terminal of the sequence considered is S (Ser).

The estimated half-life is: 1.9 hours (mammalian reticulocytes, in vitro).
>20 hours (yeast, in vivo).
>10 hours (Escherichia coli, in vivo).

Instability index:

The instability index (II) is computed to be 43.57

This classifies the protein as unstable.

Aliphatic index: 102.89

Grand average of hydropathicity (GRAVY): -0.227
