## supplemental tables for "Non-lethal exposure to H_2_O_2_ boosts bacterial survival and evolvability against oxidative stress": table S6.pdf

Table S6. Transcripts differentially expressed (±2 log2) in the fraction of small of RNA <200 nt)during the decay of hydrogen peroxide response.

| Gene | Log2 fold change | P-value | Description |
|---|---|---|---|
| oxyS | 3.73 | 8.24E-15 | OxyS sRNA regulates genes in response to H2O2 |
| rbsA | 3.12 | 2.83E-09 | Ribose import ATP-binding protein RbsA |
| rplW | 3.10 | 8.37E-08 | 50S ribosomal protein L23 |
| cyoB | 2.76 | 1.23E-06 | Cytochrome bo(3) ubiquinol oxidase subunit 1 |
| rplB | 2.45 | 4.73E-16 | 50S ribosomal protein L2 |
| rpsJ | 2.42 | 3.43E-10 | 30S ribosomal protein S10 |
| rplC | 2.24 | 5.83E-14 | 50S ribosomal protein L3 |
| fruB | 2.20 | 3.59E-04 | Multiphosphoryl transfer protein |
| leuZ | 2.11 | 1.15E-08 | Leucine tRNA(GAG) 4 |
| ptsG | 2.10 | 3.73E-10 | PTS system glucose-specific EIICB component |
| rpsC | 2.08 | 4.01E-07 | 30S ribosomal protein S3 |
| rplX | 2.06 | 1.30E-05 | 50S ribosomal protein L24 |
| rpsE | 2.03 | 2.05E-08 | 30S ribosomal protein S5 |
| raiA | -2.01 | 5.78E-12 | Ribosome-associated inhibitor A |
| dcuA | -2.05 | 1.51E-03 | Anaerobic C4-dicarboxylate transporter DcuA |
| deoA | -2.11 | 1.43E-04 | Thymidine phosphorylase |
| adhE | -2.20 | 1.15E-06 | Aldehyde-alcohol dehydrogenase |
| gatD | -2.22 | 1.41E-03 | Galactitol-1-phosphate 5-dehydrogenase |
| dnaK | -2.56 | 7.42E-05 | Chaperone protein DnaK |
| yhjX | -2.60 | 1.19E-04 | Uncharacterized MFS-type transporter YhjX |
| glnA | -2.62 | 1.12E-04 | Glutamine synthetase |
| treB | -2.66 | 5.70E-07 | PTS system trehalose-specific EIIBC component |
| glpT | -2.67 | 1.24E-06 | Glycerol-3-phosphate transporter |
| clpA | -2.76 | 1.36E-05 | ATP-dependent Clp protease ATP-binding subunit ClpA |
| glpA | -2.78 | 6.94E-05 | Anaerobic glycerol-3-phosphate dehydrogenase subunit A |
| nanA | -2.80 | 6.12E-05 | N-acetylneuraminate lyase |
| gatC | -2.88 | 2.15E-09 | NA |
| glpQ | -2.98 | 6.54E-09 | Glycerophosphoryl diester phosphodiesterase |
| glpK | -2.99 | 1.79E-08 | Glycerol kinase |
| glpB | -3.02 | 1.28E-05 | Anaerobic glycerol-3-phosphate dehydrogenase subunit B |
| glpD | -3.06 | 6.99E-09 | Aerobic glycerol-3-phosphate dehydrogenase |
| dtpB | -3.08 | 2.06E-06 | Dipeptide and tripeptide permease B |
| rbbA | -3.26 | 1.73E-06 | Ribosome-associated ATPase |
| malM | -3.61 | 5.57E-08 | Maltose operon periplasmic protein |
| tnaA | -3.78 | 9.07E-11 | Tryptophanase |
| yjiY | -4.20 | 4.75E-15 | Inner membrane protein YjiY |
