## supplemental tables for "Non-lethal exposure to H_2_O_2_ boosts bacterial survival and evolvability against oxidative stress": table S7.pdf

Table S7. Transcripts differentially expressed (± 2.5 log2) in the fraction of large RNA (>200 nt) during the decay of hydrogen peroxide response.

| Gene | Log2 fold change | P-value | Description |
|---|---|---|---|
| oxyS | 4.02 | 2.34E-10 | OxyS sRNA regulates genes in response to H2O2 |
| ptsG | 3.92 | 4.87E-96 | PTS system glucose-specific EIICB component |
| rbsA | 3.90 | 3.78E-73 | Ribose import ATP-binding protein RbsA |
| fruB | 3.66 | 2.13E-105 | Multiphosphoryl transfer protein |
| rplV | 3.48 | 7.18E-139 | 50S ribosomal protein L22 |
| rpsQ | 3.47 | 5.52E-110 | 30S ribosomal protein S17 |
| yihL | 3.42 | 2.45E-10 | Uncharacterized HTH-type transcriptional regulator YihL |
| rpsC | 3.26 | 4.84E-164 | 30S ribosomal protein S3 |
| rpsS | 3.22 | 1.96E-154 | 30S ribosomal protein S19 |
| rplP | 3.21 | 9.71E-115 | 50S ribosomal protein L16 |
| rplW | 3.13 | 5.30E-173 | 50S ribosomal protein L23 |
| rpmC | 3.10 | 3.41E-40 | 50S ribosomal protein L29 |
| cyoD | 3.05 | 6.09E-19 | Cytochrome bo(3) ubiquinol oxidase subunit 4 |
| rplB | 2.99 | 2.76E-161 | 50S ribosomal protein L2 |
| cyoC | 2.98 | 3.55E-32 | Cytochrome bo(3) ubiquinol oxidase subunit 3 |
| rpsA | 2.74 | 1.13E-167 | 30S ribosomal protein S1 |
| rplO | 2.71 | 1.56E-73 | 50S ribosomal protein L15 |
| rplQ | 2.71 | 4.27E-113 | 50S ribosomal protein L17 |
| rplD | 2.69 | 1.14E-102 | 50S ribosomal protein L4 |
| sdaB | 2.66 | 4.27E-29 | L-serine dehydratase 2 |
| fecB | 2.62 | 2.78E-06 | Fe(3+) dicitrate-binding periplasmic protein |
| fruK | 2.62 | 1.21E-36 | 1-phosphofructokinase |
| rpmD | 2.61 | 1.67E-33 | 50S ribosomal protein L30 |
| cyoB | 2.60 | 2.70E-42 | Cytochrome bo(3) ubiquinol oxidase subunit 1 |
| rbsD | 2.59 | 2.25E-95 | D-ribose pyranase |
| secY | 2.53 | 1.04E-157 | Protein translocase subunit SecY |
| rpsE | 2.52 | 9.45E-50 | 30S ribosomal protein S5 |
| rplC | 2.51 | 6.52E-149 | 50S ribosomal protein L3 |
| yicG | -2.53 | 3.42E-10 | UPF0126 inner membrane protein YicG |
| glgX | -2.55 | 1.50E-15 | Glycogen debranching enzyme |
| yhaM | -2.55 | 2.25E-07 | NA |
| treA | -2.56 | 4.75E-07 | Periplasmic trehalase |
| htpG | -2.56 | 2.54E-32 | Chaperone protein HtpG |
| frdB | -2.58 | 4.04E-04 | Fumarate reductase iron-sulfur subunit |
| ftnB | -2.59 | 3.02E-05 | Bacterial non-heme ferritin-like protein |
| aceA | -2.60 | 1.93E-13 | Isocitrate lyase |
| srlE | -2.61 | 1.06E-07 | Glucitol/sorbitol-specific phosphotransferase enzyme IIB component |
| glnA | -2.61 | 3.74E-60 | Glutamine synthetase |
| ygfF | -2.65 | 2.67E-04 | Uncharacterized oxidoreductase YgfF |
| nanE | -2.67 | 1.78E-05 | Putative N-acetylmannosamine-6-phosphate 2-epimerase |
| malM | -2.68 | 6.31E-15 | Maltose operon periplasmic protein |
| flgG | -2.71 | 6.60E-04 | Flagellar basal-body rod protein FlgG |
| glpD | -2.74 | 8.49E-13 | Aerobic glycerol-3-phosphate dehydrogenase |
| tnaC | -2.75 | 2.86E-25 | Tryptophanase leader peptide |
| yhhJ | -2.75 | 1.31E-08 | Inner membrane transport permease YhhJ |
| glnQ | -2.75 | 1.71E-09 | Glutamine transport ATP-binding protein GlnQ |
| rbbA | -2.76 | 3.25E-10 | Ribosome-associated ATPase |
| glnL | -2.78 | 2.37E-09 | Nitrogen regulation protein NR(II) |
| tdcE | -2.78 | 7.91E-09 | PFL-like enzyme TdcE |

| Gene | Log2 fold change | P-value | Description |
|---|---|---|---|
| glnH | -2.80 | 4.66E-40 | Glutamine-binding periplasmic protein |
| nanT | -2.82 | 6.98E-08 | Putative sialic acid transporter |
| dppF | -2.86 | 1.36E-05 | Dipeptide transport ATP-binding protein DppF |
| gatD | -2.88 | 2.42E-05 | Galactitol-1-phosphate 5-dehydrogenase |
| malG | -2.91 | 1.58E-06 | Maltose transport system permease protein MalG |
| srlD | -3.00 | 2.31E-16 | Sorbitol-6-phosphate 2-dehydrogenase |
| preT | -3.01 | 2.93E-13 | NAD-dependent dihydropyrimidine dehydrogenase subunit PreT |
| glnP | -3.02 | 1.34E-11 | Glutamine transport system permease protein GlnP |
| hybA | -3.06 | 5.68E-08 | Hydrogenase-2 operon protein HybA |
| dppD | -3.08 | 6.15E-05 | Dipeptide transport ATP-binding protein DppD |
| glgA | -3.09 | 2.36E-18 | Glycogen synthase |
| preA | -3.11 | 9.50E-18 | NAD-dependent dihydropyrimidine dehydrogenase subunit PreA |
| nanA | -3.18 | 7.54E-53 | N-acetylneuraminate lyase |
| glgP | -3.25 | 8.54E-28 | Glycogen phosphorylase |
| nrdD | -3.27 | 3.28E-07 | Anaerobic ribonucleoside-triphosphate reductase |
| glgC | -3.32 | 5.62E-28 | Glucose-1-phosphate adenylyltransferase |
| nanK | -3.47 | 4.04E-10 | N-acetylmannosamine kinase |
| glpK | -3.47 | 7.37E-98 | Glycerol kinase |
| srlB | -3.53 | 1.96E-05 | Glucitol/sorbitol-specific phosphotransferase enzyme IIA component |
| lldD | -3.60 | 1.00E-09 | L-lactate dehydrogenase |
| yhcH | -3.85 | 2.78E-10 | Uncharacterized protein YhcH |
| yjiY | -3.87 | 2.85E-24 | Inner membrane protein YjiY |
| dsdX | -3.88 | 1.18E-11 | DsdX permease |
| clpB | -3.91 | 9.50E-71 | Chaperone protein ClpB |
| glpQ | -4.05 | 1.21E-59 | Glycerophosphoryl diester phosphodiesterase |
| yhjX | -4.09 | 3.34E-44 | Uncharacterized MFS-type transporter YhjX |
| tdcD | -4.44 | 7.60E-07 | Propionate kinase |
| cadB | -4.67 | 1.67E-09 | Probable cadaverine/lysine antiporter |
| tnaA | -4.80 | 8.72E-29 | Tryptophanase |
| glpB | -4.94 | 9.23E-20 | Anaerobic glycerol-3-phosphate dehydrogenase subunit B |
| tnaB | -4.99 | 3.17E-13 | Low affinity tryptophan permease |
| dsdA | -5.06 | 3.05E-36 | D-serine dehydratase |
| cadA | -5.34 | 8.06E-13 | Lysine decarboxylase. inducible |
| glpC | -5.97 | 1.59E-16 | Anaerobic glycerol-3-phosphate dehydrogenase subunit C |
