## supplemental tables for "Non-lethal exposure to H_2_O_2_ boosts bacterial survival and evolvability against oxidative stress": Table S8.pdf

| Gene | Gene function | Type | Result | Expression | Std. Error | 95% C.I. | P(H1) |
|---|---|---|---|---|---|---|---|
| ***asnA*** | Asparagine synthase A | REF | | 2.3 | | | |
| ***metL*** | Bifunctional aspartokinase/homoserine dehydrogenase 2 | REF | | -2.3 | | | |
| ***mntH*** | Mn transport | TRG | UP | 55.5 | 51.567 - 59.843 | 49.843 - 61.254 | 0.000 |
| ***sufA*** | Scaffold protein for assembly of iron-sulfur clusters; facilitates delivery to target proteins; Fe-S transfer protein | TRG | UP | 18.2 | 14.303 - 21.615 | 13.185 - 25.838 | 0.000 |
| ***fur*** | Ferric uptake regulation | TRG | NDE | -1.1 | 0.845 - 0.942 | 0.832 - 0.959 | 0.090 |
| ***ahpC*** | Alkyl hydroperoxide reductase, subunit C; reduced by the AhpF subunit. protects aerobic, phosphate-starved cells from oxidative damage | TRG | UP | 7.3 | 6.616 - 8.502 | 6.507 - 8.808 | 0.000 |
| ***yaaA*** | Peroxide resistance protein, lowers intracellular iron | TRG | UP | 2.2 | 1.954 - 2.526 | 1.862 - 2.610 | 0.000 |
| ***oxyR*** | Oxidative and nitrosative stress transcriptional regulator | TRG | NDE | -4.9 | 0.185 - 0.222 | 0.179 - 0.238 | 0.079 |
| ***katG*** | Catalase-peroxidase HPI, heme b-containing; hydroperoxidase I. KatG protects aerobic, phosphate-starved cells from oxidative damage | TRG | UP | 25.7 | 19.066 - 33.678 | 17.159 - 42.580 | 0.011 |
| ***ahpF*** | Alkyl hydroperoxide reductase, subunit F; NAD(P)H:peroxiredoxin oxidoreductase, reduces AhpC; contains one FAD/monomer | TRG | UP | 5.6 | 4.739 - 6.291 | 4.489 - 7.307 | 0.011 |
| ***dps*** | Stress-induced Fe-binding and storage protein; forms biocrystals with DNA; copper homeostatsis. Dps confers starvation-induced resistance to H2O2. | TRG | UP | 3.1 | 2.676 - 3.484 | 2.530 - 3.915 | 0.021 |
