## supplemental tables for "Non-lethal exposure to H_2_O_2_ boosts bacterial survival and evolvability against oxidative stress": Table S9.pdf

Table S9. Strains and plasmids used in this work and their relevant phenotypes

| Strain | Relevant phenotype | Reference |
|---|---|---|
| *E. coli MG1655 WT* | $F^{-}$ $\lambda^{-}$ | Lab stock |
| *E. coli MG1655 oxyR::scar* | *$F^{-}$ $\lambda^{-}$ oxyR::scar* | this work |
| *E. coli MG1655 katG::scar* | *$F^{-}$ $\lambda^{-}$ katG::scar* | this work |
| *E. coli MG1655 ahpF::scar* | *$F^{-}$ $\lambda^{-}$ ahpF::scar* | this work |
| *E. coli MG1655 ahpF::scar katG::scar* | *$F^{-}$ $\lambda^{-}$ ahpF::scar katG::scar* | this work |
| *E. coli MG1655 grxA::scar* | *$F^{-}$ $\lambda^{-}$ grxA::scar* | this work |
| *E. coli MG1655 ghrA::scar* | *$F^{-}$ $\lambda^{-}$ ghrA::scar* | this work |
| *E. coli MG1655 lipA::scar* | *$F^{-}$ $\lambda^{-}$ lipA::scar* | this work |
| *E. coli MG1655 rdgB::scar* | *$F^{-}$ $\lambda^{-}$ rdgB::scar* | this work |
| *E. coli MG1655 yaaA::scar* | *$F^{-}$ $\lambda^{-}$ yaaA::scar* | this work |
| *E. coli MG1655 recA::scar* | *$F^{-}$ $\lambda^{-}$ recA::scar* | this work |
| *E. coli MG1655 fimE (Δ1 bp, 248 nt)* | *fimE* Δ1 bp, position 248 | this work |
| *E. coli MG1655 insB1→flhD Δ1 bp* | Δ1 bp, position 1978504 | this work |
| *E. coli MG1655 insB1→flhD Δ10 bp* | Δ10 bp, position 1978493 | this work |
| *E. coli MG1655 oxyS::Cm* | *$F^{-}$ $\lambda^{-}$ oxyS::Cm* | 27 |
| *E. coli BW25113 oxyR::Kan* | *oxyR::Kan* | 57 |
| *E. coli BW25113 katG::Kan* | *katG::Kan* | 57 |
| *E. coli BW25113 ahpF::Kan* | *ahpF::Kan* | 57 |
| *E. coli BW25113 grxA::Kan* | *grxA::Kan* | 57 |
| *E. coli BW25113 ghrA::Kan* | *ghrA::Kan* | 57 |
| *E. coli BW25113 lipA::Kan* | *lipA::Kan* | 57 |
| *E. coli BW25113 rdgB::Kan* | *rdgB::Kan* | 57 |
| *E. coli BW25113 yaaA::Kan* | *yaaA::Kan* | 57 |
| *E. coli BW25113 recA::Kan* | *recA::Kan* | 57 |
| pCA24N | Cloning vector, Cm-R | 36 |
| pCA24N-*fimE* | pCA24N carrying *fimE*, Cm-R | 36 |
| pBAD24 | Cloning vector, Amp-R | 80 |
| pVN15 | pBAD24 carrying *flhDC* operon, Amp-R | 80 |
