## supplemental tables for "Non-lethal exposure to H_2_O_2_ boosts bacterial survival and evolvability against oxidative stress": Table S10.pdf

Table S10. Primers used for amplification gene mutants from the Keio collection which PCR products were used to transfer the mutations to the *E. coli* MG1655.

| Gene | Forward primer (5'--->3') | Reverse primer (5'--->3') |
|---|---|---|
| *oxyR* | ACTCTCGAAACGGGCAGTG | GGTCAGGCGATTATGGAACAG |
| *katG* | GATCTCAACTATCGCATCCGTG | CACAACCAGGCCACTGAT |
| *ahpF* | ACCTAATTCTTCGGGTGCTG | GGTGCTGGTTGTGCGTTAA |
| *ghrA* | CACCGACCACGGATTTGTTATG | GCATCTTCCATATCCGGGCC |
| *grxA* | CCTCTGCAAAGTGAGCCTTC | CACCCTGTTCGATGCTCATTAT |
| *lipA* | ACCGCTTTGGCTGCTTTC | CATAAAGAGTGACGTGGCGA |
| *rdgB* | CTGGTGAAGTTTCTCGGTAAGC | CGTACCACCGCCAATAAAGA |
| *gltD* | GCGCGGTGAAGAGATTCTG | GGAAAGGTCAAACGCTCATGC |
| *yaaA* | CCCGGTGTTTGATCCATTGC | CAGGCAAACATCACCGCT |
| *recA* | GTCGTCAGGCTACTGCGTATG | GTCGCAGTTCTTGCTCACTG |
