## supplemental tables for "Non-lethal exposure to H_2_O_2_ boosts bacterial survival and evolvability against oxidative stress": Table S11.pdf

Table S11. Primers used for relative gene expression quantification by real time PCR (qPCR). RNA was extracted after a treatment of *E. coli* MG1655 with $H_2O_2$ 0.1 mM during 30 minutes.

| Gene | EcoGene ID | Forward primer (5'--->3') | Reverse primer (5'--->3') |
|---|---|---|---|
| *ahpC* | EG11384 | GCAGCACAGTACGTAGCTTCTCA | GCCAGAGTTGCTTCACCTTCTT |
| *ahpF* | EG11385 | CTGAACCTGATGAGCGTA | GACCAAACTCTTTCCCGT |
| *dps* | EG11415 | GTACATGAAATGCTGGATGG | TGCTGTTGATAACTTGAGTG |
| *fur* | EG10359 | CAGCAACATCACCACGATCAC | TGGAATCATCACTAAATTCGATAACC |
| *katG* | EG10511 | CAAATGCCCGTTCCATC | TTAACAGGTCAACACGAAGT |
| *mntH* | EG14157 | GTTATTCCGCCACCAAATG | TAGCCATCATCGCCAGA |
| *oxyR* | EG10681 | GCCAGCCGACGCTTAGC | AACATCACGCCCAGCTCATC |
| *sufA* | EG11378 | TTGATGGCACGGAAGTC | TGGGCTTTAGGGTTGTG |
| *yaaA* | EG10011 | GAAACCTTCAGCGAAGACGATT | CCATACAAGCCGGAAAGCA |
| *metL* | EG10590 | TGAGCAGGATGAAGAGTCGTTG | CCGTGGCTGGCGAAATCAAG |
| *asnA* | EG10091 | CTGGGCGGGAATTAAAGCAACC | CTCCTGGCTGTGTACGAAGTGG |
